# Endometrial cyclin A2 deficiency is associated with human female infertility and is recapitulated in a conditional knockout mouse model

**DOI:** 10.1101/2023.06.16.545284

**Authors:** Fatimah Aljubran, Katelyn Shumacher, Amanda Graham, Sumedha Gunewardena, Courtney Marsh, Michael Lydic, Kristin Holoch, Warren B. Nothnick

## Abstract

Proper action of the female sex steroids, 17β-estradiol (E2) and progesterone (P4) on endometrium is essential for fertility. Beyond its role in regulating the cell cycle, cyclin A2 (CCNA2) also mediates E2 and P4 signaling *in vitro*, but a potential role in modulating steroid action for proper endometrial tissue development and function is unknown. To fill this gap in our knowledge, we examined human endometrial tissue from fertile and infertile women for CCNA2 expression and correlated this with pregnancy outcome. Functional assessment of CCNA2 was validated *in vivo* using a conditional Ccna2 uterine deficient mouse model while *in vitro* function was assessed using human cell culture models. We found that CCNA2 expression was significantly reduced in endometrial tissue, specifically the stromal cells, from women undergoing in vitro fertilization who failed to achieve pregnancy. Conditional deletion of Ccna2 from moue uterine tissue recapitulated the inability to achieve successful pregnancy which appears to be due to alterations in the process of decidualization, which was confirmed using in vitro models. From these studies, we conclude that CCNA2 expression during the proliferative/regenerative stage of the menstrual cycle acts as a safeguard allowing for proper steroid responsiveness, decidualization and pregnancy. When CCNA2 expression levels are insufficient there is impaired endometrial responsiveness, aberrant decidualization and loss of pregnancy.

**Graphical Abstract:** 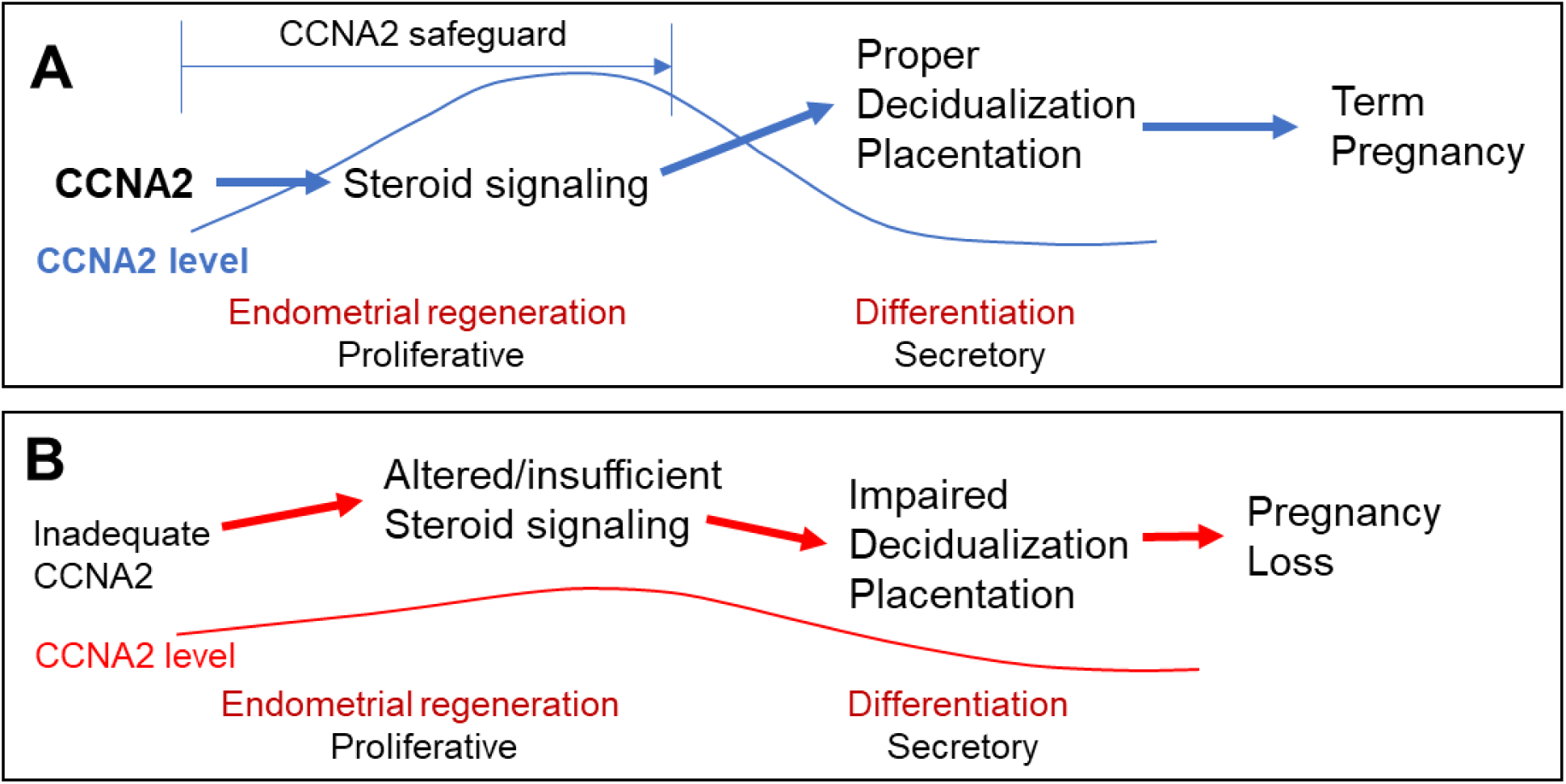

## Introduction

Infertility is a significant public health problem that affects 19% of reproductive age women in the US (1). Common categories of female infertility include ovulatory disorders, tubal occlusion, and uterine disorders (2). However, the spectrum of female infertility extends to include pregnancy disorders that leads to the loss of the embryo such as implantation failure, miscarriages or pregnancy loss. Implantation failure or the absence of clinical pregnancy accounts for 50% of infertility cases (3). On the other hand, pregnancy loss is defined as the loss of the fetus before 20 weeks of gestation which impacts 10-15% of pregnant women (4). Growing evidence suggests that abnormal uterine function plays a major role in implantation failure and pregnancy loss (3, 5). For instance, defective decidualization has been implicated in poor pregnancy outcomes such as recurrent pregnancy loss and preeclampsia (6–8).

The uterus provides a niche for the embryo to implant and grow through an intricate system that connects the uterine endometrium with the embryonic tissue known as the placenta. This process requires dynamic remodeling of the uterine lining to accommodate the implanting embryo. Moreover, cyclical remodeling of the endometrium is greatly dependent upon steroid hormone action mediated by their receptors.

Decidualization is the transformation of endometrial stromal fibroblasts into specialized secretory, decidual cells and is essential for embryo implantation, placentation and successful pregnancy. Not surprising, defective stromal cell decidualization has been linked to pregnancy complications in women including pregnancy loss and miscarriage (7). Unfortunately, our understanding of how defective stromal cell decidualization leads to pregnancy loss is poor. Studies which have examined decidualization and pregnancy establishment have primarily utilized samples from the secretory stage of the menstrual cycle and/or the “window of implantation.” While this approach is logical and outcomes have undoubtedly provided important information on mediators and mechanisms required for proper decidualization and pregnancy, assessment of only the latter half of a menstrual cycle may limit a full understanding of the events which are necessary for proper decidualization. The necessity for proper endometrial preconditioning, tissue regeneration and proliferation which occurs in the first half of a menstrual cycle for successful pregnancy has been proposed (9,10) but few, if any, studies have mechanistically explored this postulate.

Cyclins are regulatory proteins that initiates the entry and progression of the cell cycle through the activation of cyclin-dependent kinases (CDK). Growing evidence suggests that cyclins and CDKs elicit non-canonical functions in diverse cellular models (11). Cyclins have been shown to regulate steroid receptor action in CDK dependent and independent manner (12–16). Cyclin A2 (CCNA2) is predominantly expressed during the S phase of the cell cycle and interacts with CDK1 and CDK2 to activate downstream targets and induce the progression of the cell cycle (11). CDKs are proline-directed kinases that phosphorylates proteins containing Ser/Thr-Pro motifs such as the steroid receptors and their coactivators (12,13). Interestingly, cyclin A2/CDK2 complex has been shown to enhance estrogen receptor alpha (ERα) transcriptional activity in vitro through phosphorylation (13). Likewise, cyclin A2/CDK2 phosphorylates progesterone receptor (PGR) and steroid receptor coactivator 1 (SRC1) leading to the potentiation of PGR transcriptional activity in vitro (12,14–16).

Although the endometrium is a steroid responsive tissue, the potential role of CCNA2 in endometrial function has not been explored. We hypothesized that reduced expression of uterine CCNA2 may be associated with aberrant endometrial steroid signaling leading to reproductive dysfunction and infertility. To better understand the potential role of CCNA2 in endometrial signaling, we created a conditional knockout mouse model using progesterone receptor cre *(Pgr^cre^*) recombinase to mediate cyclin A2 (*Ccna2*) deletion in the female reproductive tract. The present study reveals that CCNA2 deficiency *in vivo* and *in vitro* is associated with impaired endometrial responsiveness to steroid signaling leading to pregnancy loss in utero which may be due to defective decidualization and/or placentation.

## Results

### Endometrial CCNA2 expression is reduced in women who fail to achieve IVF-assisted pregnancy

Endometrial biopsies were obtained from each subject during the early proliferative (EP) or late proliferative (LP) stages of an ovarian stimulation protocol as defined under “Materials and Methods.” Subjects undergoing in vitro fertilization (IVF) for advanced maternal age/diminished ovarian reserve, endometriosis, tubal factor infertility, recurrent pregnancy loss (RPL), unexplained infertility or male factor infertility (Table 1) provided biopsies. Pregnancy outcomes were assessed 12 to 14 days after a future frozen embryo transfer by measuring serum β-hCG. A positive pregnancy was defined as a β-hCG value of > 5 IU, while < 5 IU was considered a negative pregnancy outcome (Table 1).

**Table 1.** Patient demographics.

| TABLE 1. Patient infertility classification and pregnancy outcomes. |  |  |  |
| --- | --- | --- | --- |
| Study group | Sample size (n) | Pregnant | Not pregnant |
| <u>Early proliferative</u> |  |  |  |
| Advanced maternal age | 6 | 3 | 3 |
| Tubal factor infertility | 1 | 1 | 0 |
| Recurrent pregnancy loss | 11 | 5 | 6 |
| Unexplained infertility | 4 | 3 | 1 |
| Male factor infertility | 6 | 4 | 2 |
| TOTAL | 28 | 16 | 12 |
| <u>Late Proliferative</u> |  |  |  |
| Advanced maternal age | 4 | 3 | 1 |
| Endometriosis | 2 | 1 | 1 |
| Tubal factor infertility | 2 | 0 | 2 |
| Recurrent pregnancy loss | 3 | 1 | 2 |
| Unexplained infertility | 2 | 2 | 0 |
| Male factor infertility | 8 | 5 | 3 |
| TOTAL | 21 | 12 | 9 |
| EP + LP TOTAL | 49 | 28 | 21 |

In EP specimens from subjects who achieved a positive pregnancy, CCNA2 expression was predominantly within the stroma, showing strong nuclear staining compared to specimens from subjects who had a negative pregnancy test (Figure 1A). Assessment of H-Scores revealed that stromal cell expression of CCNA2 was significantly higher in endometrial biopsies from women who achieved pregnancy compared to those that did not (Figure 1B). Further, in EP specimens, stromal cell expression was greater compared to epithelium within each group (Figure 1B). Assessment of *CCNA2* transcript in serial sections from the same biopsies used for IHC localization revealed no difference in the level of mRNA expression between study groups (Figure 1C). A similar pattern of localization and expression was observed in endometrial biopsies collected during the LP time point (Figure 1D and 1E), with CCNA2 expression being predominantly within the nucleus of stromal cells. However, unlike EP specimens, LP specimens expressed not only significantly greater CCNA2 levels in the stromal cells of subjects who achieved pregnancy compared to those who did not (Figure 1E), but epithelial cell expression was also significantly greater in these biopsies (Figure 1E). Also, in contrast to EP specimens, *CCNA2* transcript expression was significantly greater in LP biopsies from subjects who achieved pregnancy (Figure 1F). To verify that these elevated levels of transcript expression were not due to an increase in epithelial cell content in these biopsies, we assessed cytokeratin 18 (*KRT18*) in these same RNA samples as previously described (17) and found no significant difference in ct values (± SEM) between groups (LP Pregnant = 6.79 ± 0.42 compared to 6.93 ± 0.43 for LP Not pregnant; p = 0.824).

**Figure 1.**
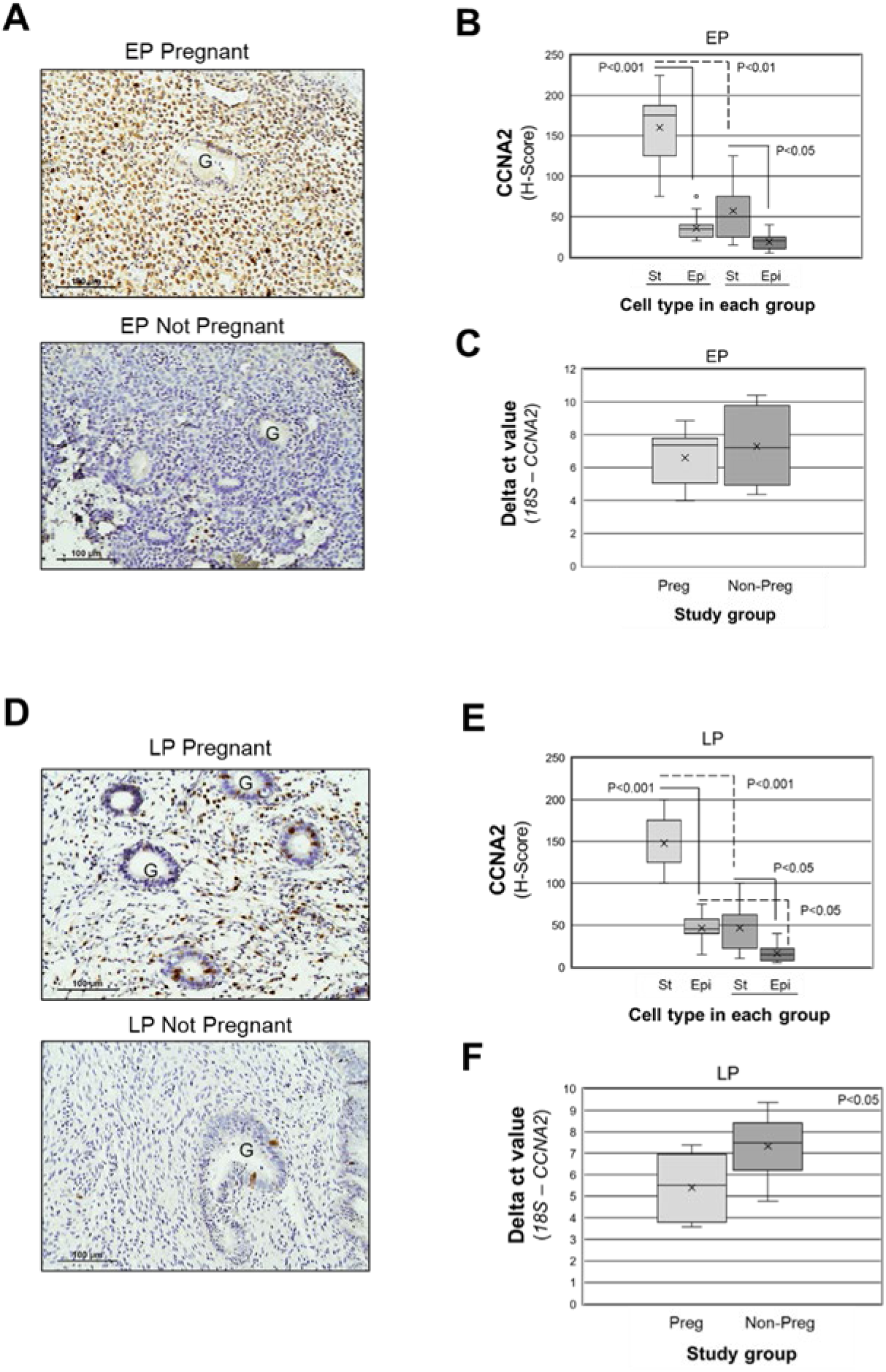
Cyclin A2 localization in early and late proliferative endometrial biopsies from women who achieved or failed to achieve ART pregnancy. A) Endometrial biopsies were obtained from women who achieved pregnancy (Pregnant) and those that did not (Not Pregnant) during the early (EP) or late (LP) proliferative stage of a stimulation cycle. Scale bar = 100 uM. G = endometrial gland. H-scores were calculated for CCNA2 expression in early (B) and late (D) proliferative biopsies. qRT-PCR was performed for CCNA2 on sections from the same blocks that IHC was performed and expressed as delta ct values (C) and fold change in CCNA2 expression (F). H-Score data were analyzed by one-way ANOVA and post-hoc analysis within cycle stage among the four groups with p-values indicated. Fold change in CCNA2 mRNA (C, F) was analyzed by unpaired t-test. n=16 EP Pregnant, n=12 EP Not Pregnant, n=12 LP Pregnant, n=9 LP Not Pregnant.

Because CCNA2 plays a critical role in cellular growth and proliferation, we aimed to assess proliferation in EP and LP endometrial biopsies of women that achieved pregnancy compared to women that did not. Immunostaining of Ki67 did not differ between women that achieved pregnancy and women that failed to achieve pregnancy (Supplemental Figure 1). However, during the late proliferative stage, proliferation index increased in endometrial stroma compared to the epithelium in both groups, but only reaching statistical significance in those in women that achieved pregnancy (Supplemental Figure 1). Because ERα and PGR mediate steroid hormones action in endometrial tissue and due to the potential role of CCNA2 in mediating these effects (12–16), we aimed to assess the expression level of ERα and PGR among women that achieved pregnancy and women that did not. In EP specimens, ERα expression was significantly greater in both epithelial and stromal cells from subjects who achieved pregnancy compared to those who did not (Supplemental Figure 2). In contrast, in LP specimens, ERα expression was greater in epithelial cells compared to stromal cells in both pregnant and not pregnant subjects. PGR expression was not significantly different in EP specimens for either epithelial or stromal cells between groups (Supplemental Figure 3). In contrast, PGR expression was significantly lower in stromal cells compared to epithelial cells from non-pregnant subjects but did not differ within cell type between study groups (Supplemental Figure 3). Collectively, these data suggested that reduced levels of CCNA2 expression in endometrial tissue from women who failed to achieve pregnancy was not associated with altered levels of Ki67, ERα, or PGR expression when compared to levels expressed in endometrial tissue from women who achieved pregnancy.

### Infertility is recapitulated in a conditional uterine knockout mouse model of Ccna2

Data from our human study suggested an inverse relationship between endometrial CCNA2 expression and pregnancy outcome. To determine if reduction/loss of CCNA2 expression functionally contributes to the ability to establish and maintain pregnancy, we developed a mouse model in which *Ccna2* was conditionally deleted from the female reproductive tract (*Ccna2^d/d^*) using Pgr^Cre^ mice to delete *Ccna2* expression within the uterus. Female mice which expressed *Ccna2* (*Ccna2^fl/fl^*) or had the gene deleted (*Ccna2^d/d^*) were mated with wild-type (C57BL/6) males of proven fertility. Females of both genotypes mated (as evidenced by a copulatory plug which is considered day post coitum [dpc] 0.5), but only *Ccna2^fl/fl^* gave birth to pups (Figure 2A). Necropsy of *Ccna2^d/d^* females at dpc 19.5 revealed the presence of resorbed fetuses (Figure 2B) in all *Ccna2^d/d^* females.

**Figure 2.**
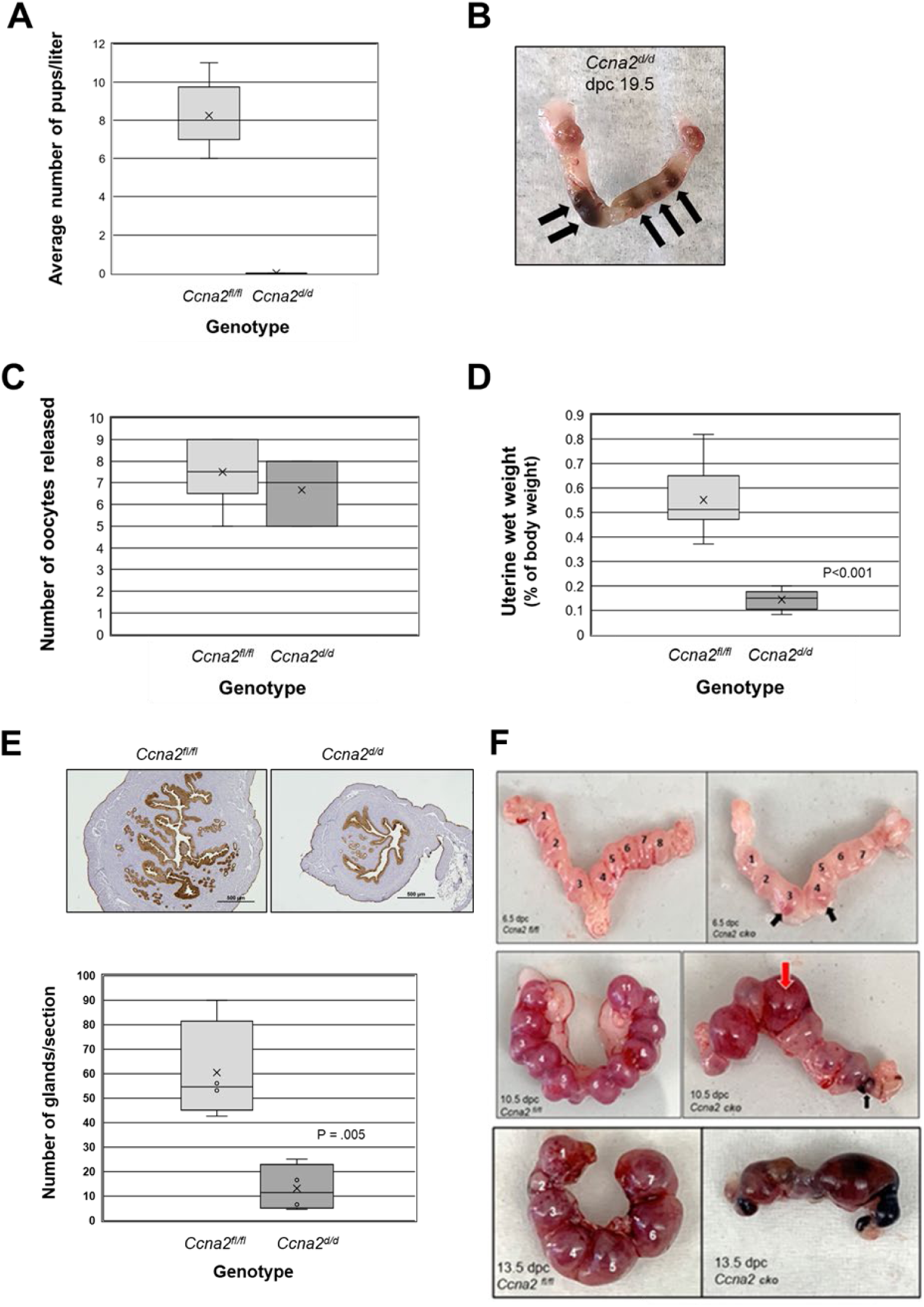
Deletion of uterine Ccna2 leads to pregnancy loss and uterine alterations. (A) Female mice in which Ccna2 is deleted from the uterus fail to give birth to offspring. At term (dpc 19.5), Ccna2 deficient mice (*Ccna2^d/d^*) exhibit fetal resorption (B). *Ccna2^d/d^*released a similar number of oocytes at dpc 0.5 (C) but had significantly smaller uteri (D) which were characterized by fewer endometrial glands (E; scale bar = 500 µm). Assessment of implantation sites at dpc 6.5, 10.5 and 13.5 (F) revealed signs of hemorrhagic implantation sites and resorption of implantation sites between dpc 6.5 (black arrows; top panel) and dpc 10.5 (middle panel; red arrows = inflammation, black arrow = resorption) in Ccna2 knockout (Ccna2 cko) mice. N = 4 – 6 mice/genotype per observation.

To begin to evaluate where in the reproductive process fetal loss may occur, we mated a separate cohort of *Ccna2^fl/fl^* and *Ccna2^d/d^*females with wild-type males of proven fertility and sacrificed females at dpc 0.5 (peri-ovulatory period), 6.5 (post embryo implantation, decidualization), 10.5 (placentation) and 13.5 (post placentation). In the dpc 0.5 groups, we first assessed ovulation rate to verify that a similar number of oocytes were released. As depicted in Figure 2C, mice of both genotypes released an average of approximately 7 oocytes. Upon harvesting the female reproductive tract to assess ovulated oocytes, it became evident that uteri from the *Ccna2^d/d^* females were considerably smaller and lacked vascularity compared to *Ccna2^fl/fl^*uteri. Assessment of uterine wet weight revealed that *Ccna2^d/d^* uteri weighted approximately one-fifth of that of *Ccna2^fl/fl^* females (Figure 2D) and were characterized by significantly fewer endometrial glands compared to *Ccna2^fl/fl^* uteri (Figure 2E). These alterations in the uterus could not be attributed to insufficient systemic levels of estradiol-17β (E2) or altered patterns of localization or levels of estradiol receptor-α (ERα) expression. Serum E2 levels were similar between genotypes (Supplemental Figure 4A) as was ERα protein localization with similar levels of expression in endometrial epithelial glands and lumen as well as stromal cells (Supplemental Figure 4B) at dpc 0.5.

We next assessed dpc 6.5 implantation sites, a time when embryos are implanted and decidualization nears completion. Mice of both genotypes exhibited a similar number of embryo implantation sites (numbered in Figure 2F; top panel), but those implantation sites in *Ccna2^d/d^* females began to exhibit signs of hemorrhage (black arrows highlight in Figure 2F; top panel), which became more apparent at mid-term pregnancy (dpc 10.5; Figure 2F; middle panel; red arrow indicates inflamed implantation site, black arrow indicates resorbed implantation site) with fetal loss apparent at dpc 13.5 (Figure 2F; bottom panel). Loss of pregnancies in the Ccna2 null mice at dpc 10.5 were not associated with insufficient serum P4 or prolactin levels in the *Ccna2^d/d^* mice as levels were actually higher compared to *Ccna2^fl/fl^* counterparts (Supplemental Figure 5).

### Loss of uterine Ccna2 is associated with an altered matrisome

To begin to assess the mechanisms by which *Ccna2* deficiency prior to embryo implantation may contribute to pregnancy loss, we performed bulk RNA-seq on dpc 0.5 uteri from mice of both genotypes to characterize the uterine transcriptome prior to the embryonic decidual signal. A total of 18,270 gene were identified in uterine tissue at dpc 0.5 (Figure 3A and Figure 3B). Using a fold change (FC) of ≥1.5, a p-value of <0.05 and a false discovery rate (FDR) of <0.1, 63 genes were significantly up-regulated (Supplemental Table 1) and 20 were significantly down-regulated (Supplemental Table 2) in dpc 0.5 uteri from *Ccna2^d/d^* mice. Pathway (Figure 3C) and process (Figure 3D) enrichment analysis revealed the top-level Gene Ontology biological processes and top 20 clusters with their representative enriched terms. Interestingly, *Ccna2^d/d^* uterine tissue exhibited significant altered expression of key components of the matrisome including extracellular matrix proteins, proteases and cytokines. Focusing on the top differentially expressed genes (Supplemental Tables 1 and 2) using cutoff values of > 2-fold change between genotypes, p<0.05 and FDR < 0.1, we validated two of the most significantly up-regulated and two of the most-significantly down-regulated transcripts. These up-regulated transcripts were identified as tenascin (*Tnc*) and cellular communication network factor 2 (*Ccn2*) which is also known as connective tissue growth factor [*Ctgf*]), while the down-regulated transcripts were identified as *Ovgp1* [oviductal glycoprotein 1] and Kruppel like factor 15 [*Klf15*]. Interestingly all four of these transcripts have been reported to be associated with embryo implantation and decidualization (18–31). All four genes displayed a similar fold change when comparing bulk RNA-seq data with that of qRT-PCR (Figure 3E). Interestingly, the majority of the significantly down-regulated genes (Supplemental Table 2) were determined to be epithelial cell enriched which may reaffirm the necessity of glandular epithelial-derived factors for proper endometrial decidualization in both mouse (32–34) and humans (35).

**Figure 3.**
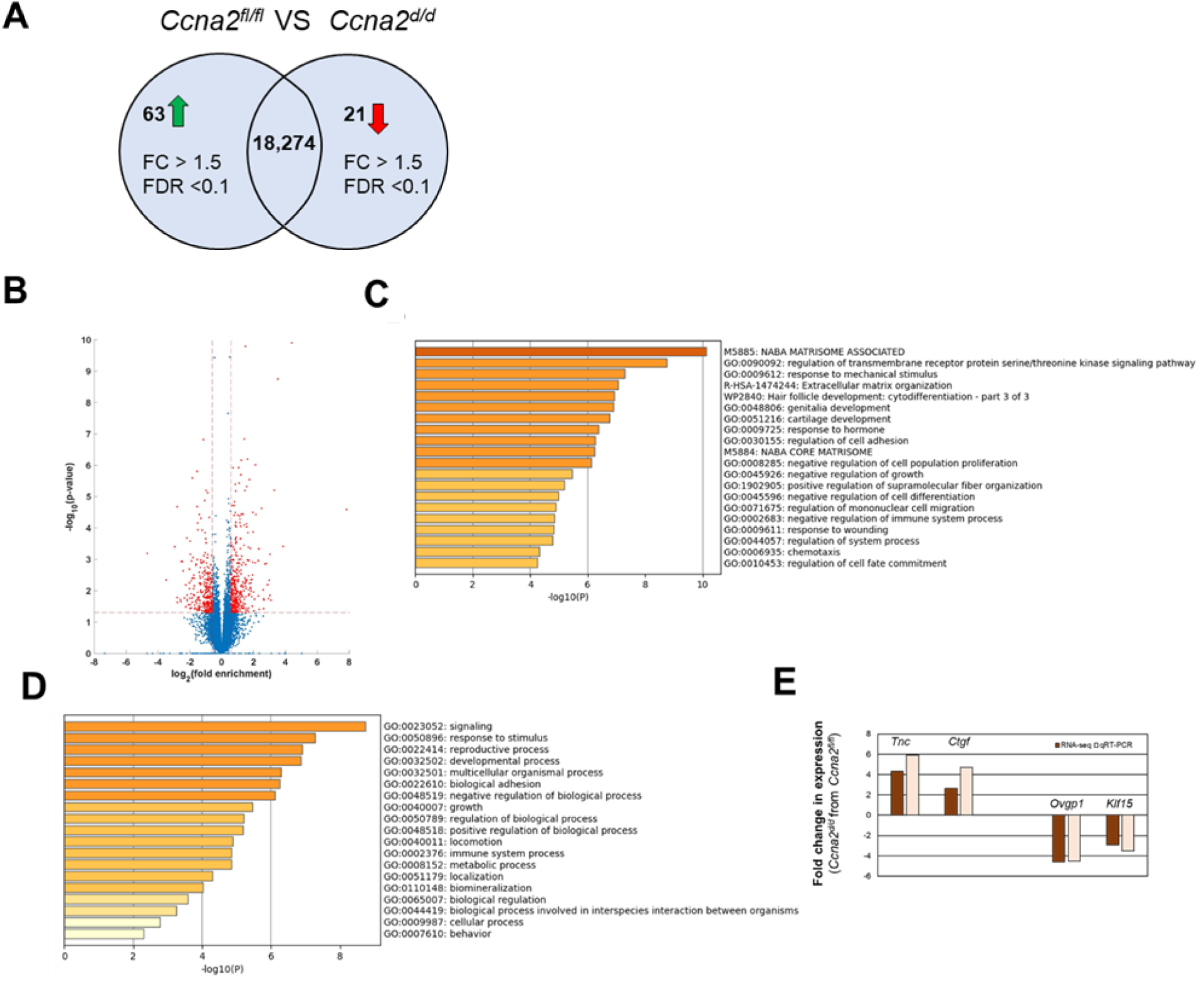
Bulk RNA-sequencing analysis in dpc 0.5 uteri. A) Venn diagram and B) Volcano plot shows the number and distribution of differentially expressed genes in dpc 0.5 uteri of *Ccna2^d/d^* mice(p-value <0.05). (C) Pathway and (D) process analysis for the differentially expressed (≥1.5 fold change, p<0.05, q<0.1) genes between genotypes in dpc 0.5 uterine tissue. (E) fold change in two of the top up-regulated and down-regulated genes by RNA-seq analysis and validation by qRT-PCR. n=4 mice/genotype used to generate the Volcano Plot and n=6 mice/genotype for qRT-PCR validation studies.

### Loss of uterine Ccna2 is associated with impaired estradiol-17β action and steroid signaling

To determine if the lower uterine wet weight characteristic of *Ccna2^d/d^* mice (Figure 2D) could be attributed to altered steroid action, female mice of both genotypes were ovariectomized and treated with E2. Uterine wet weight of *Ccna2^d/d^*mice was significantly lower compared to *Ccna2^fl/fl^* in all treatment groups (Figure 4A). Compared to the control, untreated (0h) group, *Ccna2^fl/fl^*mice treated with E2 gained uterine wet weight at 6h and 24h which was statistically significant at 24h within genotype (Figure 4A). Although E2 treated *Ccna2^d/d^*mice gained significant wet weight at 24h compared to 0h, this increase was significantly lower compared to *Ccna2^fl/fl^* group at 24h (Figure 4A) and uteri appeared less vascular and smaller from both gross and histological assessment (Figure 4B). To evaluate hormonal responsiveness of uteri from *Ccna2^d/d^* we next assessed the expression of known E2-regulated genes which are components of the uterine matrisome from mice of both genotypes (Figures 4C to 4G).

**Figure 4.**
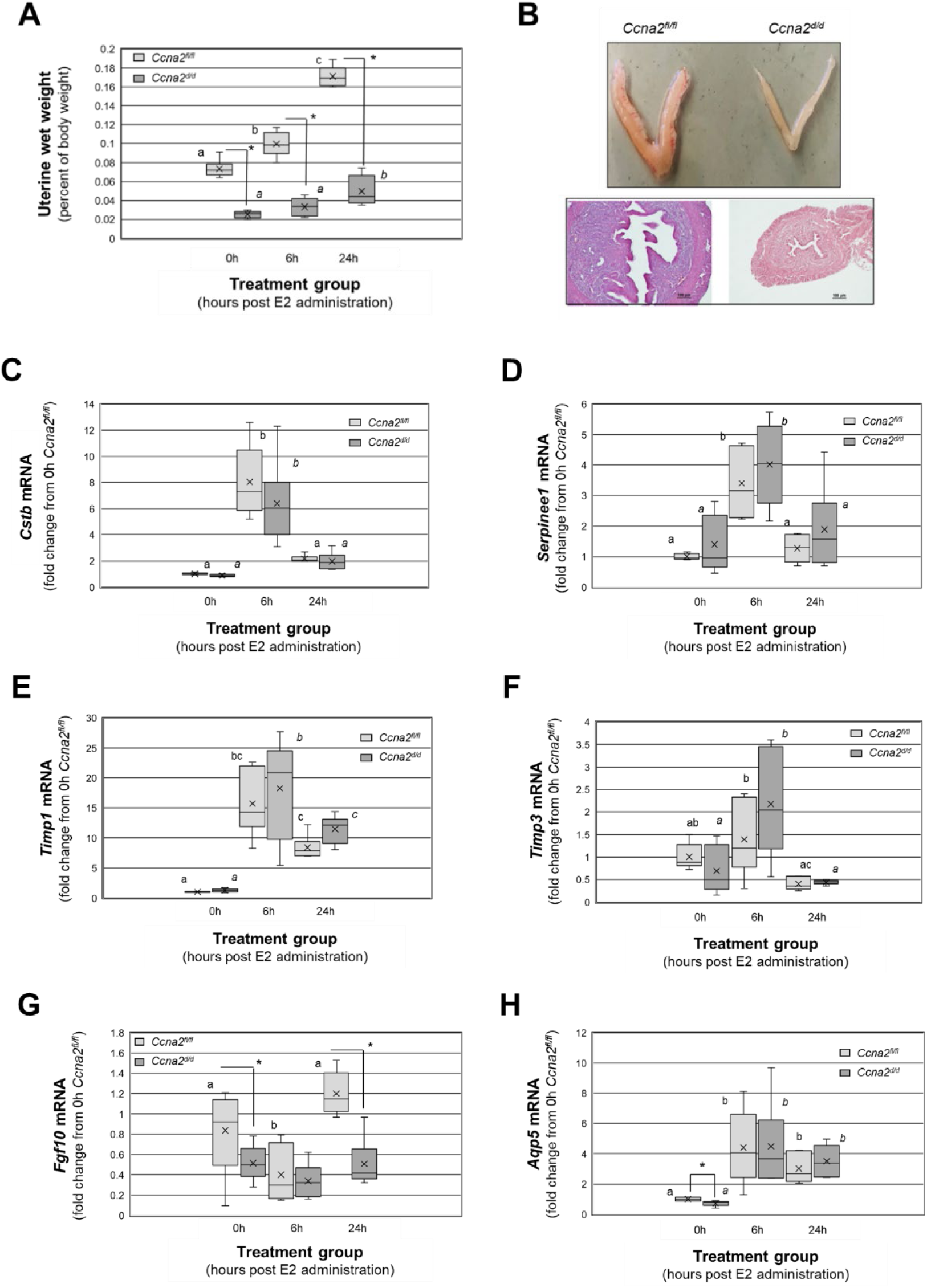
Impaired steroid action in *Ccna2^d/d^* uteri. A) Uterine wet weight at 0h (prior to treatment), 6 and 24h after E2 treatment. Different letters indicate statistical significance among the means for the *Ccna2^fl/fl^* treatment groups. There were no statistically significant differences among the means across steroid treatment within the *Ccna2^d/d^*mice. * indicates statistical significance (P<0.001) between genotypes within each treatment group (n=6 mice/group). B) Gross uterine morphology and histology in *Ccna2^fl/fl^* and *Ccna2^d/d^*mice at 24h post E2-treatment. C-E) E2 regulation of *Cstb*, *Serpine1*, *Timp1*, *Timp3*, *Fgf10* and *Aqp5* transcripts between genotypes. Different block letters indicate statistical significance among the means for the *Ccna2^fl/fl^*treatment groups while different letters in italics indicate statistically significant differences among the means in *Ccna2^d/d^* mice by one-way ANOVA. * indicates statistical significance (P<0.001) between genotypes within each treatment group (n=6 mice/group).

Transcript expression of cystatin B (*Cstb*; Figure 4C) and serpine1 (*Serpine1*; Figure 4D), which are respectively cysteine and serine protease inhibitors, was significantly increased by E2 at 6h post administration returning to control/baseline levels at 24h. There was no difference in the pattern or extent of induction of these protease inhibitors between genotypes at any of the time points, including 0h. Tissue inhibitors of metalloproteinases, *Timp1* and *Timp3* also exhibited a significant increase in expression at 6h post E2 administration (Figure 4E and Figure 4F, respectively). Estrogen increased *Timp1* expression in mice of both genotypes at 6 and 24h (Figure 4D). In contrast, *Timp3* levels only showed significantly different levels of expression in the *Ccna2^fl/fl^* uteri at 6h compared to 24h (Figure 4F). On the contrary, E2 induced a significant increase in *Timp3* expression at 6h in *Ccna2^d/d^*uteri which returned to baseline levels at 24h (Figure 4F). Like *Ctsb* and *SerpinE1*, there were no significant differences in the extent of induction of either *Timp1* or *Timp3* by E2 between genotypes (Figure 4E and Figure 4F).

Assessment of fibroblast growth factor-10 (*Fgf10*) and aquaporin-5 (*Aqp5*), which may contribute to endometrial stromal cell proliferation and uterine water imbibition, respectively, revealed slightly different patterns of expression. Compared to 0h, *Fgf10* transcript expression in *Ccna2^fl/fl^* was significantly reduced at 6h post-E2 administration and returned to baseline, control levels at 24h (Figure 4G). In contrast, *Fgf10* expression in *Ccna2^d/d^* uterine tissue did not respond to E2 treatment as expression was significantly lower compared to *Ccna2^fl/fl^* tissue levels at 0h and 24h. E2 increased *Aqp5* at both 6 and 24h post-administration and the levels of expression did not differ between genotypes at this time points (Figure 4H). At 0h, *Aqp5* expression was significantly lower in 0h uterine tissue of the in *Ccna2^d/d^* mice compared to controls (Figure 4H). Of interest was the observation that both *Fgf10* and *Aqp5* transcript expression was significantly lower in dpc 0.5 uteri from *Ccna2^d/d^* mice compared to *Ccna2^fl/fl^* mice (Figure 4G and 4H) which may suggest that these genes may be disrupted early during uterine development and may contribute to the smaller uteri detected in the *Ccna2^d/d^* mice.

### Loss of uterine Ccna2 is associated with altered decidualization in vivo

To evaluate if loss of Ccna2 prior to mating compromises the decidualization process, we assessed markers of decidualization at dpc 6.5 in mice of both genotypes. Compared to *Ccna2^fl/fl^*dpc 6.5 implantation sites, expression of *Prl3c1* (Figure 5A) and *Prl8a2* (Figure 5B) were significantly reduced in tissues from *Ccna2^d/d^* mice. Similarly, expression of the stromal cell enriched protein, *Bmp2* (Figure 5C) and *Timp3* (Figure 5D) were also significantly reduced. However, expression of *Serpine1*, which is also stromal cell enriched and associated with successful decidualization, did not change between genotypes (Figure 5E). As *Klf15* (24–26) and *Ovgp1* (21,30,31) misexpression was reported to be associated with infertility, we also assessed their expression. Both *Ovgp1* (Figure 5F) and *Klf15* (Figure 5G) expression were significantly lower in dpc 6.5 implantation sites from *Ccna2^d/d^*mice. Lastly, there were no significant differences in serum progesterone levels between genotypes (Figure 5H) at dpc 6.5, supporting the notion that pregnancy loss was not due to inadequate levels of this steroid.

**Figure 5.**
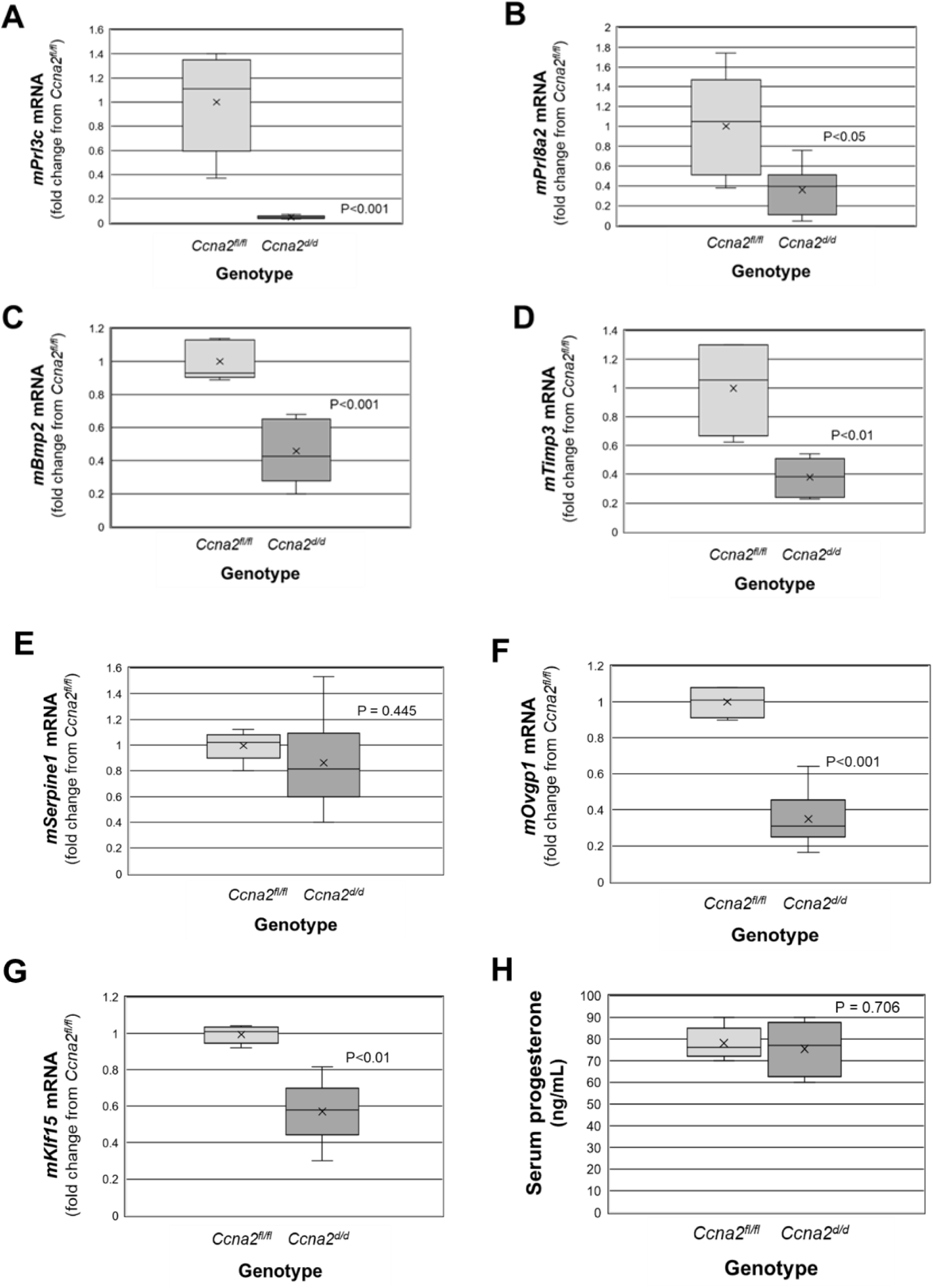
Ccna2 is essential for optimal decidualization. qRT-PCR was performed on dpc 6.5 implantation sites (1 – 2/mouse) from *Ccna2^fl/fl^* and *Ccna2^d/d^* females (n=5/genotype) for decidualization markers *Prl3c* (A), *Prl8a2* (B), *Bmp2* (C), *Timp-3* (D), *Serpine1* (E), *Ovgp1* (F), *and Klf15* (G). Serum P4 (n=5/genotype) was assessed by ELISA between genotypes (H). Data were analyzed by unpaired t-test between genotypes and p-values are presented in each figure.

### CCNA2 knockdown in vitro reduces the expression of decidualization-specific genes in t-HESC

To provide further proof of principal that CCNA2 deficiency prior to a decidualization stimulus compromises the decidualization process, we utilized the well-described *in vitro* decidualization model using the human endometrial stromal cell line, t-HESC. As the pattern of CCNA2 expression during *in vitro* decidualization is unknown, we first assess transcript expression by qRT-PCR. *CCNA2* transcript levels were highest prior to induction of *in vitro* decidualization and rapidly declined and remained low from days 2 through 10 (Figure 6A) showing an inverse relationship with the well-known markers of decidualization *PRL* (Figure 6B) and *IGFBP1* (Figure 6C). To demonstrate that reduction of CCNA2 prior to decidualization (which occurs in our *Ccna2^d/d^* mouse model) negatively impacts the decidualization process, we transfected t-HESCs with *CCNA2* siRNA or a non-targeting (NT) siRNA prior to decidualization. Twenty-four hours after transfection, we changed the media to decidualization media and harvested the cells at day 0 (prior to decidualization stimulus), day 2, and day 4 to assess the expression of decidualization markers prolactin (*PRL*) and insulin like growth factor binding protein 1 (*IGFBP1*). Knockdown of CCNA2 was confirmed in the day 2 and day 4 CCNA2-siRNA-transfected cells (Figure 6D) as was the pattern of CCNA2 expression in the NT-siRNA-transfected cells (Figure 6D). Expression of *PRL* (Figure 6E) and *IGFBP1* (Figure 6F) significantly increased in cells transfected with NT siRNA on day 2 and day 4 compared to day 0. However, the expression of *PRL* (Figure 6E) and *IGFBP1* (Figure 6F) in *CCNA2* deficient cells were significantly lower compared to control cells on day 2 and day 4. These data indicate that CCNA2 may play a critical role in endometrial steroid signaling and its expression prior to decidualization may be required for full decidualization and full term pregnancy.

**Figure 6.**
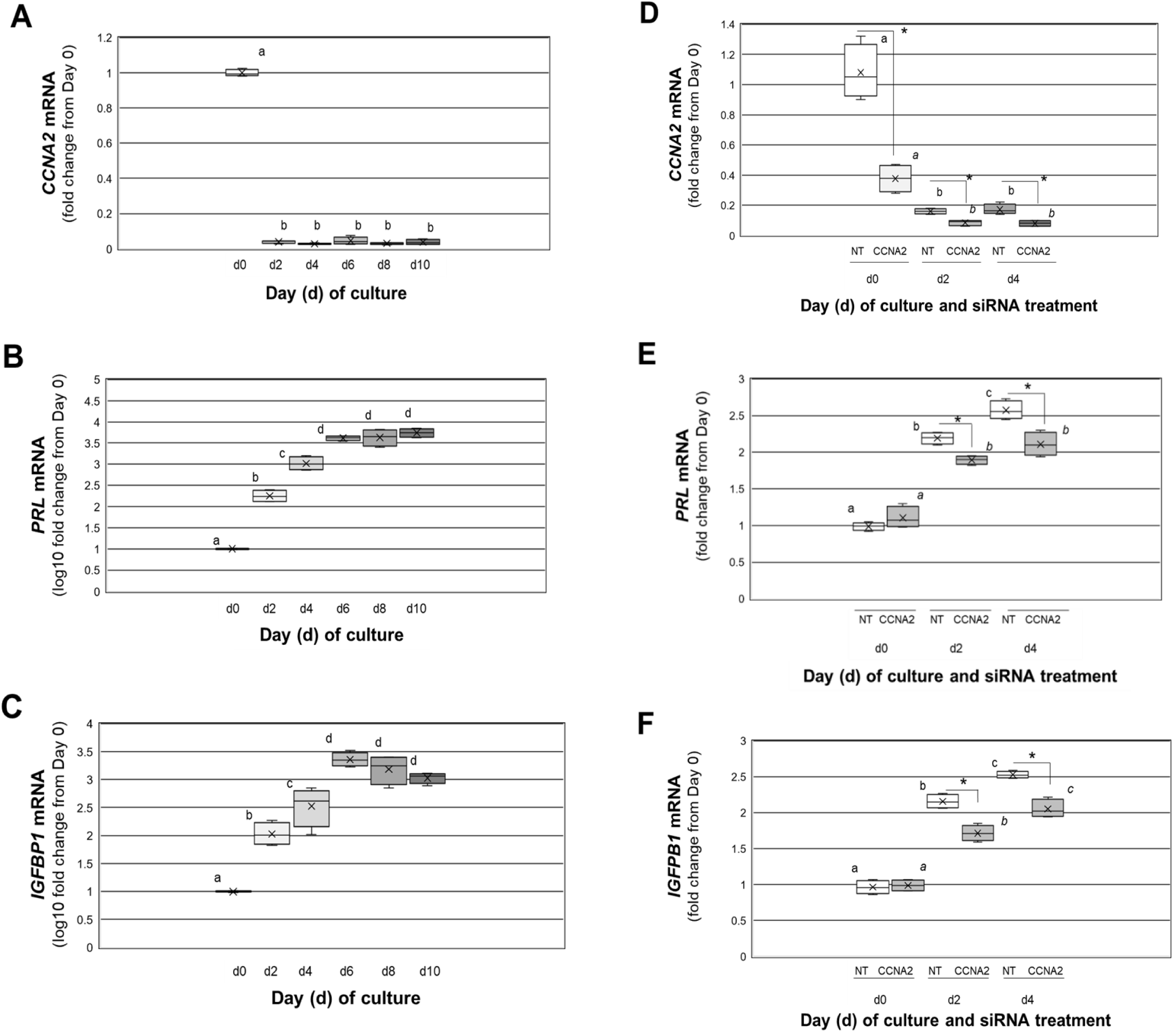
CCNA2 deficiency in a human endometrial stromal cell lines impairs decidualization. A) *CCNA2*, B) *PRL* and C) *IGFBP1* transcript expression during *in vitro* decidualization in non-transfected t-HESCs. D) *CCNA2*, E) *PRL* and F) I*GFBP1* transcript expression in NT-siRNA and CCNA2-siRNA transfected t-HESC prior to (day 0), and after a decidualization stimulus. Different block letters indicate statistical significance among the means across days of culture/treatment groups for NT-siRNA-transfected cells while different letters in italics indicate statistically significant differences among the means in the CCNA2-siRNA-transfected cells using one-way ANOVA followed by Tukey-Kramer multiple comparison tests. * indicates statistical significance (P<0.001) between transfection groups within each time point using unpaired t-tests. For Figures 6B and 6C as well as Figure 6E and 6F, values are expressed as the log10 value of the fold change from the appropriate control for each transcript. Data are from 4 separate experiments (N=4).

## Discussion

Successful pregnancy requires a suitable endometrial environment, stromal cell decidualization and placental development, all of which are dependent upon proper steroid signaling. Cyclins (CCNs) are a family of proteins that, along with their cyclin-dependent kinases (CDKs) control the progression of the cell cycle. CCNs and CDKs have been reported to exhibit non-canonical functions (11) as well as mediating E2 (12,13) and P4 (12, 14–16) signaling. Cyclins and CDKs have been examined in normal uterine tissue of humans (36–46) and mouse (47) with most of the emphasis on CCNE and CCND family members. Little information is available for CCNA2, which is one of two members of the CCNA family (CCNA1 and CCNA2). Reports on CCNA (reports did not specify if they assessed CCNA1 or CCNA2 expression) have primarily focused on endometrial carcinoma tissue, but when normal endometrium was assessed, CCNA expression was low to absent (44–46) in these tissues with any expression localized to endometrial epithelium. CCNA1is restricted to the germ line (48,49), while CCNA2 is ubiquitously expressed in all proliferating cells and is upregulated in a variety of cancers (50) including endometrial cancer (44–46). In the current study, low/deleted levels of CCNA2 in both human and mouse tissues, respectively, was not associated with major differences in cell proliferation which may be attributed to potential compensation by other cyclins, such as CCNE which was reported in the Ccna2 null mice (51).

In addition to regulating cell proliferation, CCNA2 was also demonstrated to mediate estrogen (12,13) and progesterone (12,14–16) receptor signaling. CCNA2 may modulate steroid action and act as a co-regulator. CCNA2 expression was significantly lower in biopsies from subjects who failed to achieve pregnancy but this was not associated with major differences in expression or loss of ERα or PGR expression within epithelial or stromal cells in biopsies from subjects who achieved pregnancy compared to those who did not. If CCNA2 functions as a co-regulator, then despite expression of ERα or PGR in endometrial tissue from those who fail to achieve pregnancy, E2 and/or P4 action could be diminished. This appears to be the case in our mouse model as some but not all steroid responsive transcripts and events were compromised. For example, the ability of E2 to induce uterine wet weight in ovariectomized mice was severely compromised in the *Ccna2^d/d^*mice, but E2 regulation of some target genes was less affected. More specifically, analysis of differential gene expression during the early and late E2 responses revealed that ERα responsive genes were not affected globally by the deletion of *Ccna2* in the uterus. We did not detect significant differences in the gene expression of *Cstb, Serpine1,* and *Timp1* between control and *Ccna2^d/d^* mice by qRT-PCR, however, regulation of *Fgf10* (52) was compromised as *Ccna2^d/d^* mice and exhibited significantly lower levels at 24h post E2 administration compared to *Ccna2^fl/fl^*mice.

Another novelty of the current study was the emphasis to focus on endometrial tissue specimens obtained prior to the window of implantation. The majority of studies (53–55) have focused on the endometrial environment during the secretory stage of the menstrual cycle with emphasis on the window of embryo implantation (which typically begins on day 19/20 of an idealized 28 day menstrual cycle and lasts approximately 4 to 5 days (53)). However, it is becoming apparent that events which occur during the menstrual and proliferative stages of the menstrual cycle when regeneration and proliferation of the endometrium occur may also play a role in establishing an optimal endometrial environment for successful term pregnancy. Brosens and colleagues proposed a role for menstruation in preconditioning of the uterus for successful pregnancy (9) while Bromer and associates (10) postulated that diseases associated with infertility may manifest a developmental defect during the proliferative phase of the menstrual cycle which may contribute to the difficulty to establish pregnancy. However, until recently little mechanistic studies have been conducted to support or refute these postulates. A recent study by Lv and coworkers (56) evaluated the endometrial niche of human thin endometrium in specimens obtained during the late proliferative phase of the subject’s normal menstrual cycle using single-cell RNA sequencing. In subjects with thin endometrium (defined as an endometrial thickness < 7 mm at mid-luteal phase or histories of embryo transfer cancellations in vitro fertilization procedures due to thin endometrium) the investigators reported inhibition of stromal cell proliferation in thin endometrium specimens which was associated with impaired EGF, PTN, and TWEAK signaling pathways and overactivation of the SEMA3B pathway. Further, the defective growth of the thin endometrium/decreased proliferation was associated with increased cellular senescence and collagen deposition. One of the most striking observations in our study with respect to the *Ccna2^d/d^* null mice was the hypoplastic uteri which exhibited a thin endometrium compared to *Ccna2^fl/fl^*mice in both the naturally mated dpc 0.5 groups as well as the E2-treated ovariectomized mice. However, in the study by Lv and associates (56), thin endometrium was associated with increased expression of markers of cell senescence (*p16*, *p21*) and collagen deposition (*Col4a1*). In our study, we did not detect significant differences in these genes in *Ccna2^d/d^* mice although the genes for p16 (*Cdkn2a*) and p21 (*Cdkn1a*) exhibited higher expression levels (approximately 3- and 2-fold, respectively but p-values and false discovery rate values did not meet our inclusion criteria) compared to *Ccna2^fl/fl^*mice, while *Col4a1* was not differentially expressed between genotypes. Potential explanations may be related to species differences or that fact that in the study by Lv and colleagues (56), “thin” endometrium was obtained from patients who had multiple D&C procedures while the controls did not undergo such procedure which may confound outcomes. With respect to our human specimens in our study, only 1 subject had a prior D&C procedure but this subject achieved pregnancy on a subsequent frozen embryo transfer cycle.

In assessing infertility (failed embryo implantation, early pregnancy loss), most focus has been on the window of implantation. Recent evidence suggests a suitable implantation environment may first be established prior to embryo implantation during the proliferative stage of the menstrual cycle if not earlier. Our animal model in which *Ccna2* is deleted from the uterine tissue prior to puberty would support this notion.

Similar to the observation of increased collagen/extracellular matrix (ECM) deposition in thin endometrium and infertility in humans (56), uteri from *Ccna2^d/d^* mice at dpc 0.5 exhibited alterations in expression of genes associated with the matrisome, an ensemble of genes encoding ECM proteins and ECM-associated proteins. Of these, we confirmed increased expression of tenascin C (*Tnc*) and connective tissue growth factor (*Ctgf*; also known as *Ccn2*). Both *Tnc* (46,47) and *Ctgf* (48, 49) are steroid regulated genes and may potentially play a role in recurrent pregnancy loss/altered decidualization. Of the down regulated genes, two of the most significantly decreased genes were epithelial-specific genes, *Klf15* (24–26) and *Ovgp1* (21,30,31) and like *Tnc* and *Ctgf*, both are steroid-regulated and have been associated with embryo implantation and decidualization as well. Currently, how disruption of Ccna2 expression may lead to increased expression of *Tnc* and *Ctgf* is unknown as is the mechanism for reduced expression of *Klf15* and *Ovgp1*. Considering that both *Klf15* and *Ovgp1* are epithelial-specific genes, the observation that the total number of endometrial glands are reduced in *Ccna2^d/d^* uteri may offer some explanation for their reduced expression. The necessity for endometrial glands (and their secretions) for successful pregnancy is clearly established (32–35). Despite a reduction in the number of endometrial glands, *Ccna2^d/d^* females were still capable of achieving early pregnancy but were incapable of supporting pregnancy to term. Whether the reduced number of glands in these mice contributed to later loss of pregnancy is unknown.

Perhaps the most significant finding from the current study was the observation that endometrial tissue from women who fail to achieve pregnancy after an ART-assisted cycle express significantly lower levels of CCNA2 and that conditional deletion of *Ccna2* from mouse uterine tissue results in an inability to carry a pregnancy to term, with the loss of pregnancy occurring between endometrial decidualization and placentation. The mechanism by which CCNA2 may impact decidualization prior to a decidualization stimulus remains uncertain but both our human in vitro and mouse in vivo data strongly support this postulate. Of the known transcription factors with documented roles in the decidualization process (57), only FosB proto-oncogene, AP-1 transcription factor subunit (Fosb) was differentially expressed in the dpc 0.5 uteri of *Ccna2^d/d^*mice exhibiting a 2.1-fold increase in expression compared to control counterparts. Fosb is one of the four FOS family members that can dimerize with proteins of the JUN family to form the AP-1 transcription complex (58), which in turn induces expression of key decidualization factors (59). How increased Fosb may impede decidualization is unknown but may be associated with altered cell dynamics. Over-expression of FOSB decreased the transformation phenotype of gastric cancer cell lines in vitro (60) and Fosb expression was linked to elevated levels of cyclo-oxygenase-2 (COX2) in colon adenocarcinoma HCA-7 cells (61). While COX2 plays a vital role in decidualization (62), it is tempting to speculate that premature or inappropriate temporal induction of this enzyme may lead to altered decidualization.

In summary, we report for the first time that CCNA2 levels are significantly lower in endometrial tissue obtained during controlled ovarian stimulation protocols who fail to achieve pregnancy compared to those who do. Using a conditional Ccna2 knockout mouse model in which Ccna2 is deleted from uterine tissue, we further demonstrate that loss of this cyclin results in impaired decidualization and ability to maintain term pregnancy. Future investigation to identify the mechanisms and mediators through which CCNA2 may regulate decidualization are currently underway.

## Material and Methods

### Human samples

This study was approved by the Institutional Review Board at the University of Kansas Medical Center. Our cohort study consists of women undergoing in vitro fertilization (IVF) cycles for infertility treatment. The patients were diagnosed with one of the following conditions: male factor only, tubal factor infertility, advanced maternal age, recurrent pregnancy loss, and unexplained infertility (Table 1). Endometrial samples were obtained from a non-conception cycle and pregnancy outcomes were reported as positive/negative pregnancy based upon beta-hCG levels. Biopsies were obtained during the early proliferative (cycle day 3-6; *n* = 28) and late proliferative (cycle day 11-13; *n* = 21) stages of the controlled stimulation cycle. Endometrial biopsies (50 – 100 mg of tissue) were obtained using a pipelle instrument and placed directly into 10% neutral-buffered formalin for transport to the research laboratory. Endometrial tissue was prepared for paraffin embedding and serial sectioning. Resulting tissue sections (5 µm sections) were mounted onto glass slides for immunohistochemistry.

### Immunohistochemistry (IHC) and H-score

To assess the expression and localization pattern of CCNA2, Ki67, ERα, and PR in the endometrium, IHC was performed following the protocol of the reagent supplied (Vector Laboratories Inc. Burlingame, CA) using commercially available antibodies (Supplemental Table 3). First, the slides containing the tissue section are deparaffinized with xylene then rehydrated with series of 100%, 95%, and 75% ethanol. After a washing step, slides are incubated in antigen-unmasking solution at 95°C (H-3300, Vector Laboratories Inc.) for antigen retrieval. After that, slides are incubated in 1X phosphate buffered saline (PBS) on ice, then quenched with 3% hydrogen peroxide. The VECASTAIN ABC kit (Vector labs PK-6101) is used for IHC assay. Slides are incubated with diluted normal serum provided by the kit, then primary antibody for 1 hour at room temperature. Then, slides are washes and incubated with the biotinylated secondary antibody provided by the kit. After that Avidin/Biotinylated Enzyme complex (ABC) is added. After that, ImmPACT DAB peroxidase (HRP) substrate (SK-4100, Vector Laboratories, Inc.) is used to develop a brown color. Then slides are counterstained with Hematoxylin and dehydrated with graded ethanol and xylene. Then slides are mounted and coverslip with VectaMount (H-5000, Vector Laboratories, Inc.). The expression level and localization of each target is quantified using the H-score system. Each slide was blindly evaluated at 20X field by two reviewers. The intensity of staining is scored individually as (0 = no staining, 1 = minimal staining, 2 = moderate, 3 = strong) of each cell type (stroma and glandular epithelium). The average intensity is calculated using the following equation: (H-score for each cell type = average intensity X % of cell type X 100%) as previously reported by our group (63).

### Animals

Floxed *Ccna2* mice on a C57BL/6 background (64) were crossed with *Pgr^cre^* mice (C57BL/6 background) to generate conditional knockout mice. The cre recombinase mediate *Ccna2* deletion in *Pgr* expressing tissues (*Ccna2^fl/fl^*; *Pgr^cre/+^*, referred here in as *Ccna2^d/d^)*. Floxed *Ccna2* mice were generously provided by Dr. Peter Sicinski at Dana-Farber Cancer Institute, Boston, MA, while *Pgr^cre^* mice were generously provided by Dr. John P Lydon at Baylor College of Medicine, Houston, TX. Control (*Ccna2^fl/fl^*) and *Ccna2^d/d^* female mice were sacrificed at the indicated ages. At the indicated time points of sacrifice, uterine morphology was grossly evaluated and uterine wet weight determined. Measurement of body weight and uterine wet weight are taken prior to tissue processing for endpoint analysis. The uterus was divided and preserved in RNALater solution or 10% Neutral Buffered Formalin (NBF) until processed for RNA isolation or immunohistochemistry as described below.

### Genotyping

To obtain DNA, ear punch biopsies were obtained from mice between 14 and 17 post-natal days of age. DNA was extracted and PCR reactions performed to identify the genotype of the mouse. DNA extraction was done using REDExtract-N-Amp Tissue PCR kit (XNAT, Sigma). For the PCR reaction, REDExtract-N-Amp PCR ReadyMix (R4775, Sigma) was used. CCNA2 fl/fl primers: CCNA2 loxP1 forward 5’ -CGCAGCAGAAGCTCAAGACTCGAC- 3’, CCNA2 loxP1 reverse 5’ -TCTACATCCTAATGCAATGCCTGG- 3’, and CCNA2 deleted 5’ - CACTCACACACTTAGTGTCTCTGG- 3’. PR-Cre primers: PR-Cre 1 5’ - ATGTTTAGCTGGCCCAAATG- 3’, PR-Cre 2 5’ -TATACCGATCTCCCTGGACG- 3’, and PR-WT 5’ - CCCAAAGAGACACCAGGAAG- 3’.

### Fertility assessment

Control (*Ccna2^fl/fl^*) and *Ccna2^d/d^*female mice were mated with wild-type (C57BL/6) males of proven fertility. Females were checked daily for presence of copulatory plugs to confirm mating, with the detection of a copulatory plug considered day post coitum (dpc) 0.5. Plugged females were housed separately and allowed to deliver to term. On term date, the number of pups born were recorded. In the case of no pups born, *Ccna2^d/d^* were sacrificed to assess fetal resorption.

### Ovariectomy and hormonal treatment

*Ccna2^fl/fl^* and *Ccna2^d/d^* female mice were ovariectomized following standard procedures at 2 – 3 months of age, rested 14 days and then treated with estradiol (E2; 10 µg/kg BW). Mice were sacrificed at the indicated time points, uterine tissue wet weight determined and uterine tissues were processed for either histological assessment or RNA extraction and subsequent qRT-PCR analysis.

### Evaluation of pregnancy

*Ccna2^fl/fl^* and *Ccna2^d/d^* female mice were mated with wild-type (C57BL/6) males of proven fertility as described under “Fertility assessment.” Plugged females were housed separately and sacrificed at dpc 0.5, 2.5, 6.5, 10.5, or 13.5. On dpc 0.5, the oocytes were retrieved from the oviduct by gently tearing open the oviduct with fine forceps and counted. Blood samples were collected immediately after decapitation from all mice at all dpc timepoints. Serum samples was obtained by allowing blood samples to clot at room temperature for 30 – 45 minutes followed by centrifugation at 3800 rpm at 22 C°. The supernatant (serum) was removed into clean 1.5 mL microfuge tubes and stored at −80 C° until assessed for estradiol, progesterone and/or prolactin content. Uterine samples were obtained immediately after blood collection for RNA assessment and histology.

### Assessment of endometrial gland density

To assess the number of endometrial glands, tissue sections from mice of both genotypes (at dpc 0.5) were subjected to IHC localization as described under “Immunohistochemistry (IHC) and H-score” for cytokeratin-19 (KRT19) to visualize glands. The number of glands per section were then manually counted using a Nikon microscope for 2 separate sections in one uterine horn for each mouse (N=4 mice/genotype). The average number of glands/cross section was then calculated by taking the average of the 2 sections.

### Stranded total RNA library RNA-sequencing

Stranded total RNA-sequencing was performed using the Illumina NovaSeq 6000 Sequencing System at the University of Kansas Medical Center – Genomics Core (Kansas City, KS). Quality control of the total RNA submissions was completed using the Agilent TapeStation 4200 using the RNA ScreenTape Assay kit (Agilent Technologies 5067-5576). Total RNA (500ng) was used to initiate the Illumina Stranded Total RNA Prep Ligation with Ribo-Zero Plus (Illumina 20040525) library preparation protocol. The total RNA fraction underwent a probe based ribosomal reduction using Ribo-Zero Plus probes which are hybridized to rRNA targets with subsequent enzymatic digestion to remove the over abundant rRNA from the purified total RNA. The ribosomal depleted RNA fractions were sized by fragmentation, reverse transcribed into cDNA, end repaired and ligated with the appropriate indexed adaptors using the IDT for Illumina RNA UD unique dual indexes (UDI) (Illumina 20040553) to yield strand specific RNA-seq libraries.

Following Agilent TapeStation D1000 QC validation of the library preparation, the final library quantification by qPCR was performed using the Roche Lightcycler96 with FastStart Essential DNA Green Master (Roche 06402712001) and KAPA Library Quant (Illumina) DNA Standards 1-6 (KAPA Biosystems KK4903). The RNA-Seq libraries were normalized to a 2nM concentration and pooled for multiplexed sequencing on the NovaSeq 6000.

Pooled libraries were denatured with 0.2N NaOH (0.04N final concentration) and neutralized with 400mM Tris-HCl pH 8.0. The pooled libraries were diluted to 380 pM prior to onboard clonal clustering of the patterned flow cell using the NovaSeq 6000 S1 Reagent Kit v1.5 (200 cycle) (Illumina 20028318). A 2×101 cycle sequencing profile with dual index reads is completed using the following sequence profile: Read 1 – 101 cycles x Index Read 1 – 10 cycles x Index Read 2 – 10 cycles x Read 2 – 101 cycles. Following collection, sequence data is converted from .bcl file format to fastq file format using bcl2fastq software and de-multiplexed into individual sequences for data distribution using a secure FTP site or Illumina BaseSpace for further downstream analysis.

### Progesterone and estradiol-17β enzyme-linked immunosorbent assay (ELISA)

Estradiol-17β (E2) serum levels were quantitated using a rat/mouse ELISA kit (ES180S-100; Calbiotech, El Cajon, CA) while progesterone (P4) serum levels were quantitated using a rat/mouse ELISA kit (IB79183, Immuno-Biological Laboratories, Inc. (IBL-America). Briefly, mice were sacrificed by decapitation at the indicated time points, blood collected and serum separated. Serum samples were undiluted (E2) or diluted 1:10 (P4; 10 µL serum + 90 µL diluent) and subjected to the assays following the provided protocols. After adding the stop solution to each well, the absorbance at 450 was determined for each sample using a microplate reader (FlexStation3 multi-mode microplate reader, Molecular Devices, San Jose, CA). The concentration for each sample was calculated using the standard curve of the calibrators provided. 17β-estradiol standard range: 3 – 300 pg/mL, sensitivity: 3 pg/mL, intra-assay coefficient of variation (CV) = 3.1%, and inter-assay CV = 9.9%. Progesterone ELISA standard range: 0.4 –100 ng/ml, sensitivity: 0.156 ng/ml, Intra-assay CV = 7.9%, and inter-assay CV = 9.6%.

### Prolactin enzyme-linked immunosorbent assay (ELISA)

To measure prolactin concentration in mice sera, blood was collected at the indicated time points as described above and serum prolactin (diluted 1:10) was quantitated using mouse prolactin ELISA PicoKine® kits (Boster Biological Technology, Pleasanton, CA following the provided protocol. After adding the stop solution to each well, the absorbance at 450 was determined for each sample using a microplate reader (FlexStation). The concentration for each sample was calculated using the standard curve of the calibrators provided. Standard range: 15.6 – 1,000 pg/mL, sensitivity: <10 pg/mL. Intra-assay CV = 4.6% and inter-assay CV = 5.8%.

### Cell culture

Immortalized human endometrial stromal cells (t-HESC; 65) were obtained from ATCC (CRL-4003; Manassas, VA) and were maintained with DMEM-F12 (Fisher/Corning) media supplemented with 10% Fetal Bovine Serum (non-Charcoal Stripped-FBS), 1% Penicillin/Streptomycin, and 2μl/ml Normocin. For cell transfection, t-HESC were seeded in six-well plates at a density of 350,000 cells/well and cultured in (DMED-F12) with 2% Charcoal Stripped-FBS and 1% Penicillin/Streptomycin. Lipofectamine 2000 was used to prepare the *CCNA2* or non-targeting (NT) siRNA solution. T-HESC were transfected with *CCNA2* siRNA or non-targeting siRNA prior to decidualization induction. Twenty-four hours later, media was changed to decidualization media which contained 5% horse serum, 1 μM medroxyprogesterone acetate, 10 nM E2, and 0.5 mM 8-bromoadenosine 3′,5′-cyclic monophosphate with media changes every 48 hours. t-HESC were harvested at day 0, day 2, and day 4 to assess the expression of decidualization markers prolactin (*PRL*) and insulin like growth factor binding protein 1 (*IGFBP1*).

### RNA isolation and qRT-PCR assessment

qRT-PCR was performed as previously described (17,63). Briefly, total RNA was isolated from tissue or cells using Tri-Reagent (Sigma Chemical Co., St. Louis, MO) according to recommendations of the manufacturer. Total RNA (1 µg in 20µl) was reverse transcribed using reverse transcription (RT) kits (Applied Biosystems; Foster City, CA) following the manufacturer’s protocol. Primers (Supplemental table 4) were designed for human and mouse target transcripts using Primer-Blast and synthesized by Integrated DNA Technology (IDT, Coralville, IA). Resulting material was then used for independent qRT-PCR which was carried out on an Applied Biosystems HT7900 Sequence Detector. To account for differences in starting material, human 18S rRNA primers (4333760; Thermo Fisher Scientific, Pittsburgh, PA.) were used, while for mouse tissues, ribosomal protein L13 primers (*mRpl13*; Supplemental table 4) were used. All samples were run in triplicate and the average value used in subsequent calculations. The 2-delta-delta CT method was used to calculate the fold-change values among samples as previously described by our group (17,63).

### Statistics

All data were first assessed for normal (Gaussian) distribution using the Kolmogorov-Smirnov test. Data which displayed normalcy of distribution were analyzed by one-way analysis of variance (ANOVA) within genotypes across treatments, while comparisons within time points/treatments between genotypes were made using unpaired t-tests. Post-hoc analysis was made using Student-Newman Keuls post-hoc analysis. Data which failed to display normality of distribution were analyzed by non-parametric tests (Mann-Whitney t-tests or Kruskal-Wallis test/one-way ANOVA on ranks) followed by post-hoc analysis using Dunn’s multiple comparison tests. Data are displayed as box and whisker plots unless otherwise specified. All analysis was conducted using GraphPad Prism6 (GraphPad Software, La Jolla, CA) and significance was set at P < 0.05.

## Study approval

For studies utilizing human tissues, the study was approved by the Institutional Review Board at the University of Kansas Medical Center. Written, informed consent was obtained from all study subjects prior to participation. Biopsies were obtained at the University of Kansas Health System Center for Advanced Reproductive Medicine by either a reproductive endocrinologist (CM, ML, KH) or a registered nurse. For animal studies, all experiments and procedures are approved by the Institutional Animal Care and Use Committee (IACUC) of University of Kansas Medical Center in accordance with the NIH Guide for the Care and Use of Laboratory Animals.

## Author contributions

The study was designed by WBN, FA, KS and CM, experiments were conducted by WBN, FA, KS and AG. CM, ML and KH identified subjects who satisfied the inclusion/exclusion criteria, consented and obtained biopsies. WBN, FA, KS and CM wrote the draft manuscript. WBN, FA and SG analyzed data. All authors read and approved the final manuscript.

## Acknowledgements

We thank the University of Kansas Medical Center - Genomics Core for generating the array data sets and the Histology Core for generating tissue blocks. The Genomics Core is supported by the University of Kansas - School of Medicine, the Kansas Intellectual and Developmental Disability Research Center (NIH U54 HD090216) and the Molecular Regulation of Cell Development and Differentiation - COBRE (5P20GM104936-10) while the Histology core is supported by the KIDDRC NIH U54 HD 090216 at the University of Kansas Medical Center, Kansas City, KS 66160. Gratitude is expressed to the REI nursing staff for patient consent and those patients who graciously gave consent for this project.

The research reported in this study was supported in part by the NIH Kansas INBRE grant P20GM1033418 through a collaboration with the University of Kansas Medical Center Lied Pre-Clinical Research Pilot Grant Program awarded to WBN.

## Supplemental figures

**Supplemental Figure 1.**
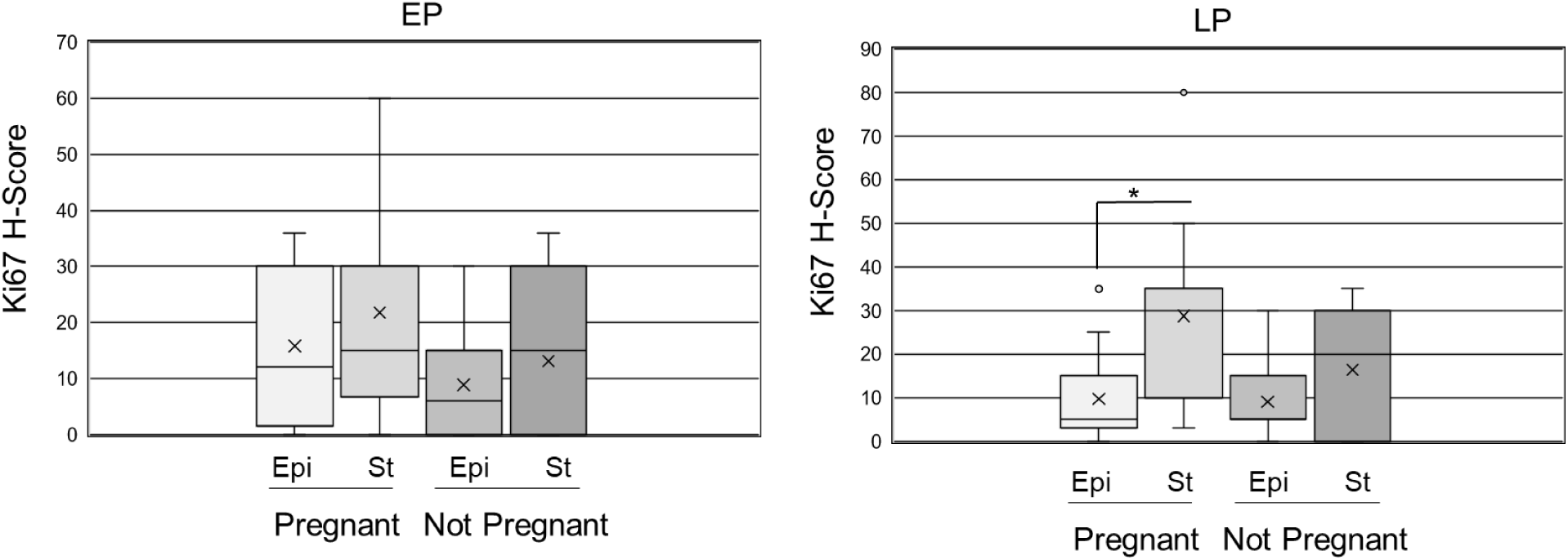
Ki67 expression in early and late proliferative endometrial biopsies from women who achieved or failed to achieve ART-assisted pregnancy. H-scores were calculated for Ki67 expression in endometrial biopsies during early proliferative (EP) and late proliferative (LP) stages of ovarian stimulation cycles. Data are presented in Box and Whisker plot of H-scores of endometrial samples in epithelial (Epi) and stromal cells (St) who either achieved a chemical pregnancy (Pregnant) or did not (Not Pregnant) for samples taken during the specified cycle days of non-transfer cycles. * indicates statistical significance (P < .05) between the stroma and epithelium within group of women who achieved pregnancy during the late proliferative stage. The (x) indicates the mean. The whisker end points indicate the highest and lowest values. H-Score data were analyzed by one-way ANOVA and post-hoc analysis within cycle stage among the four groups with p-values indicated for samples sizes of n=12 EP Pregnant, n=11 EP Not Pregnant, n=14 LP Pregnant, n=11 LP Not Pregnant.

**Supplemental Figure 2.**
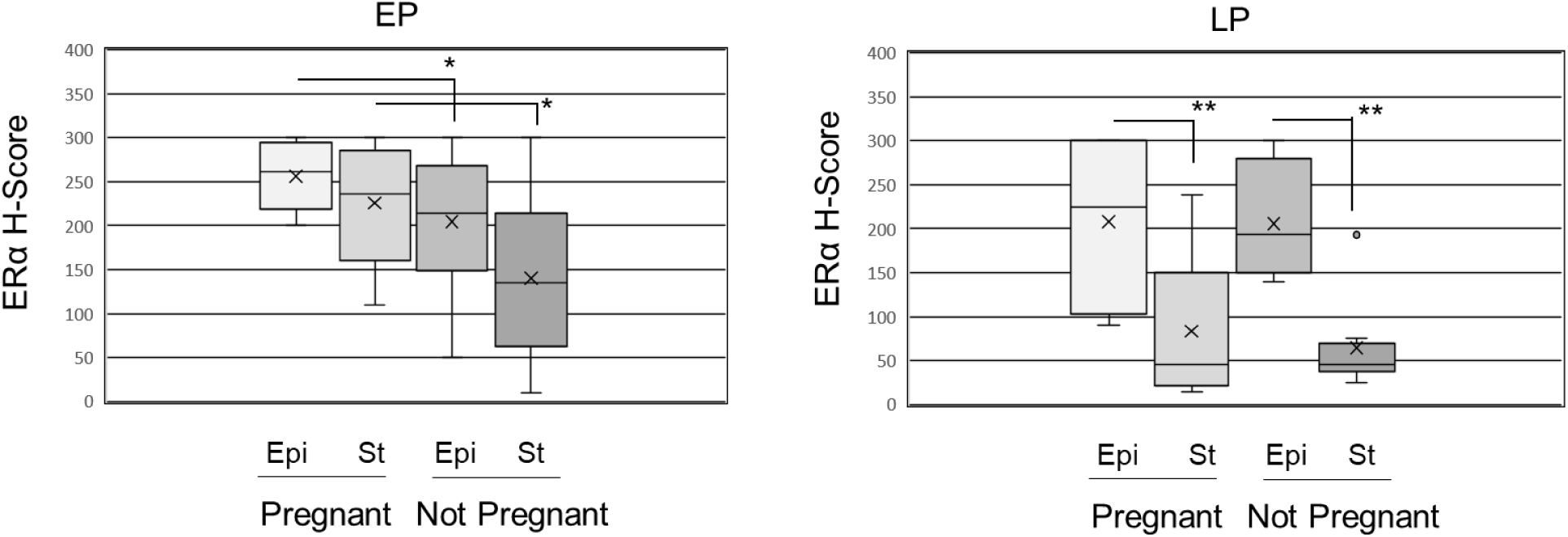
Estrogen receptor-alpha expression in early and late proliferative endometrial biopsies from women who achieved or failed to achieve ART-assisted pregnancy. H-scores were calculated for estrogen receptor-alpha (ERα) expression in endometrial biopsies during early proliferative (EP) and late proliferative (LP) stages of ovarian stimulation cycles. Data are presented in Box and Whisker plot of H-scores of endometrial samples in epithelial (Epi) and stromal cells (St) who either achieved a chemical pregnancy (Pregnant) or did not (Not Pregnant) for samples taken during the specified stage of non-transfer cycles. The (x) indicates the mean. The whisker end points indicate the highest and lowest values. H-Score data were analyzed by one-way ANOVA and post-hoc analysis within cycle stage among the four groups with p-values indicated for samples sizes of n=12 EP Pregnant, n=11 EP Not Pregnant, n=14 LP Pregnant, n=11 LP Not Pregnant. * indicates P<0.05 between epithelial and stromal cells from pregnant compared to not pregnant specimens, ** indicates P<0.01 between cell types within pregnant or not pregnant groups.

**Supplemental Figure 3.**
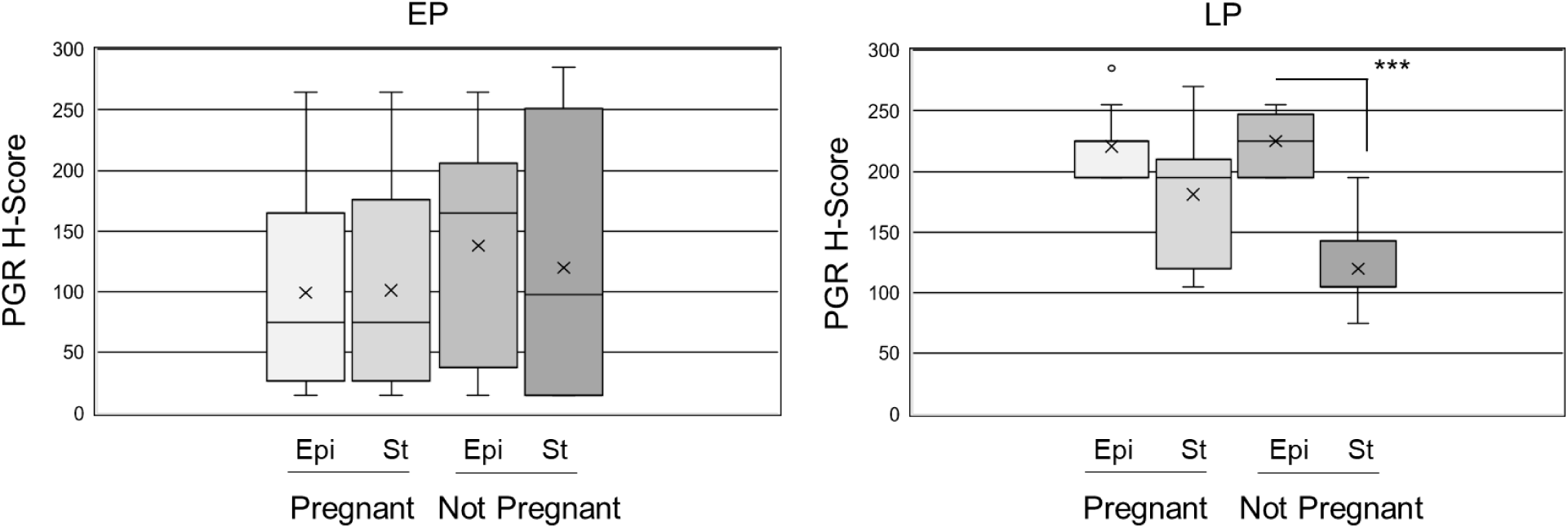
Progesterone receptor expression in early and late proliferative endometrial biopsies from women who achieved or failed to achieve ART-assisted pregnancy. H-scores were calculated for progesterone receptor A/B (PGR) expression in endometrial biopsies during early proliferative (EP) and late proliferative (LP) stages of ovarian stimulation cycles. Data are presented in Box and Whisker plot of H-scores of endometrial samples in epithelial (Epi) and stromal cells (St) who either achieved a chemical pregnancy (Pregnant) or did not (Not Pregnant) for samples taken during the specified stage of non-transfer cycles. The (x) indicates the mean. The whisker end points indicate the highest and lowest values. H-Score data were analyzed by one-way ANOVA and post-hoc analysis within cycle stage among the four groups with p-values indicated for samples sizes of n=12 EP Pregnant, n=11 EP Not Pregnant, n=14 LP Pregnant, n=11 LP Not Pregnant. *** indicates P<0.001 between epithelial and stromal cells from not pregnant LP specimens.

**Supplemental Figure 4.**
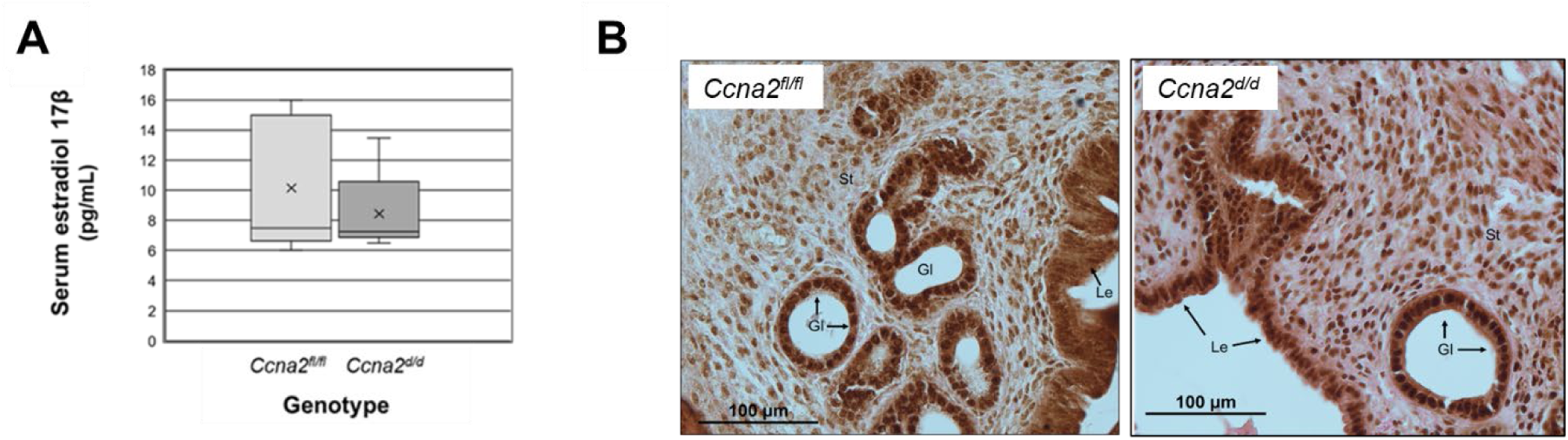
Serum estradiol levels and estradiol receptor uterine expression in *Ccna2^f/fl^*and *Ccna2^d/d^* female mice post ovulation. At dpc 0.5, trunk blood was collected and serum estradiol-17β (E2) levels were quantitated by ELISA (A) while estrogen receptor-alpha (Erα) expression was localized in uterine tissues by immunohistochemistry and sections counterstained with eosin (n=6/genotype). Brown nuclear signal represents Erα localization. Gl = glandular epithelium, Le= luminal epithelium, St = stroma; scale bar = 100 µm. Data are displayed as box and whisker plots for serum E2 (n=6/genotype) and were analyzed by unpaired t-tests. P>0.05 for comparing data between genotypes.

**Supplemental Figure 5.**
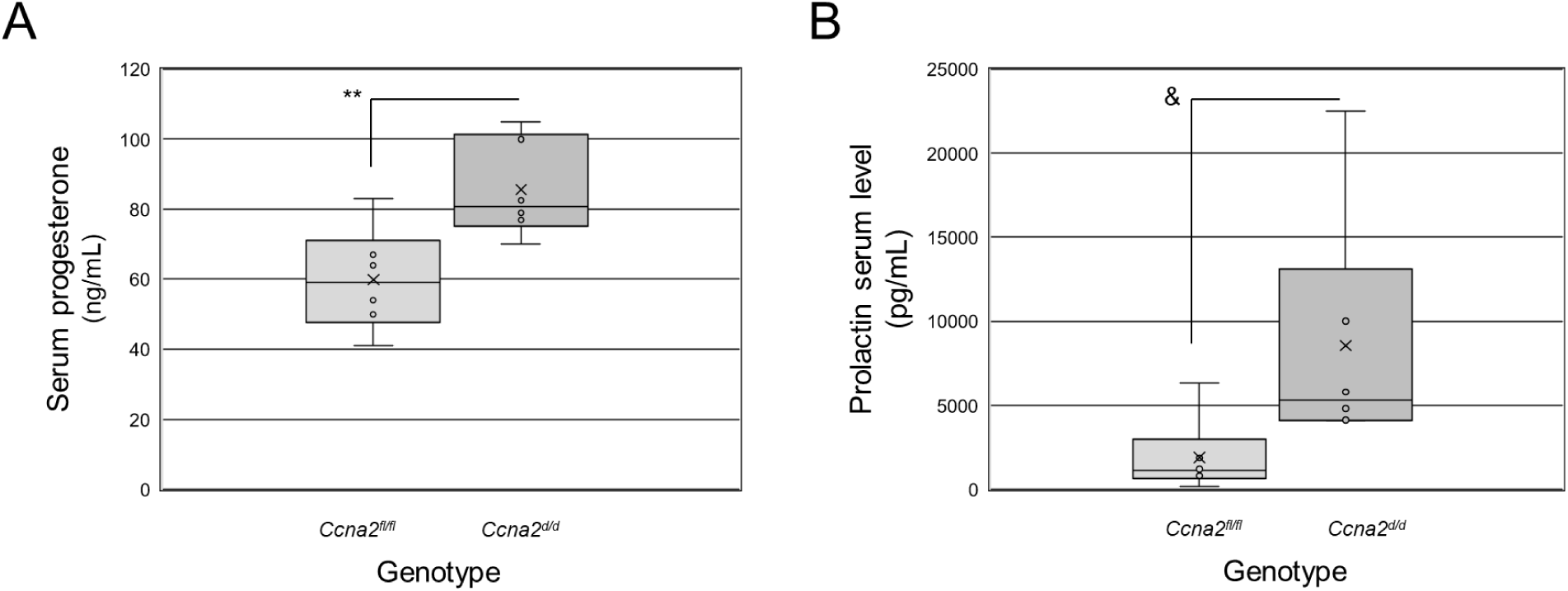
Mid-term pregnancy serum progesterone and prolactin levels in *Ccna2^f/fl^* and *Ccna2^d/d^* female mice. Serum progesterone (P4) and prolactin (Prl) were quantitated by ELISA at mid-term (dpc 10.5) pregnancy. Data are presented in Box and Whisker plots of serum P4 (A) and Prl (B) for 6 mice/genotype (n=6/genotype). Data were analyzed by unpaired t-tests for each analyte. ** indicates P<0.01; & indicates P = 0.07.

---

**Supplemental Table 1.**
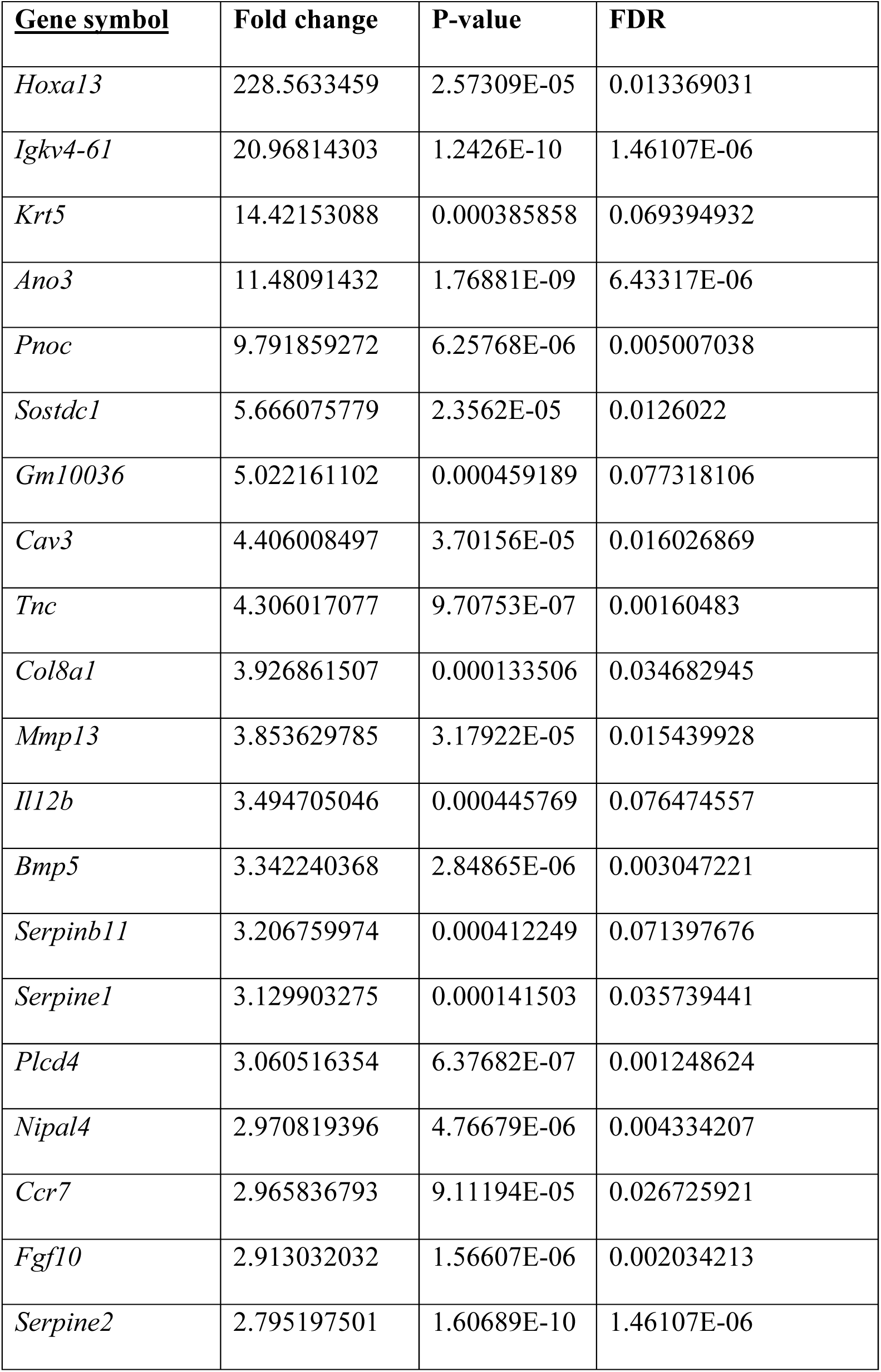
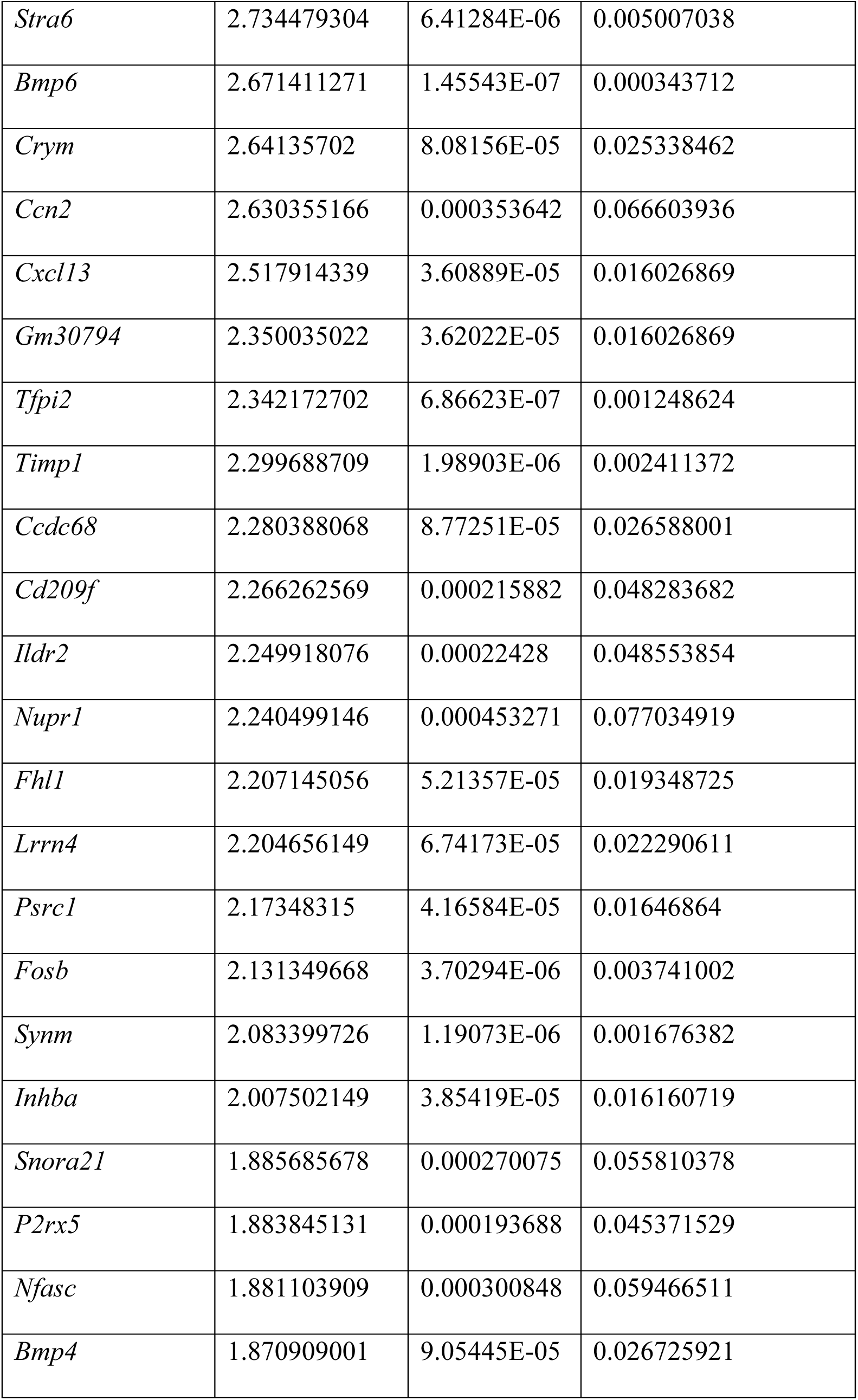
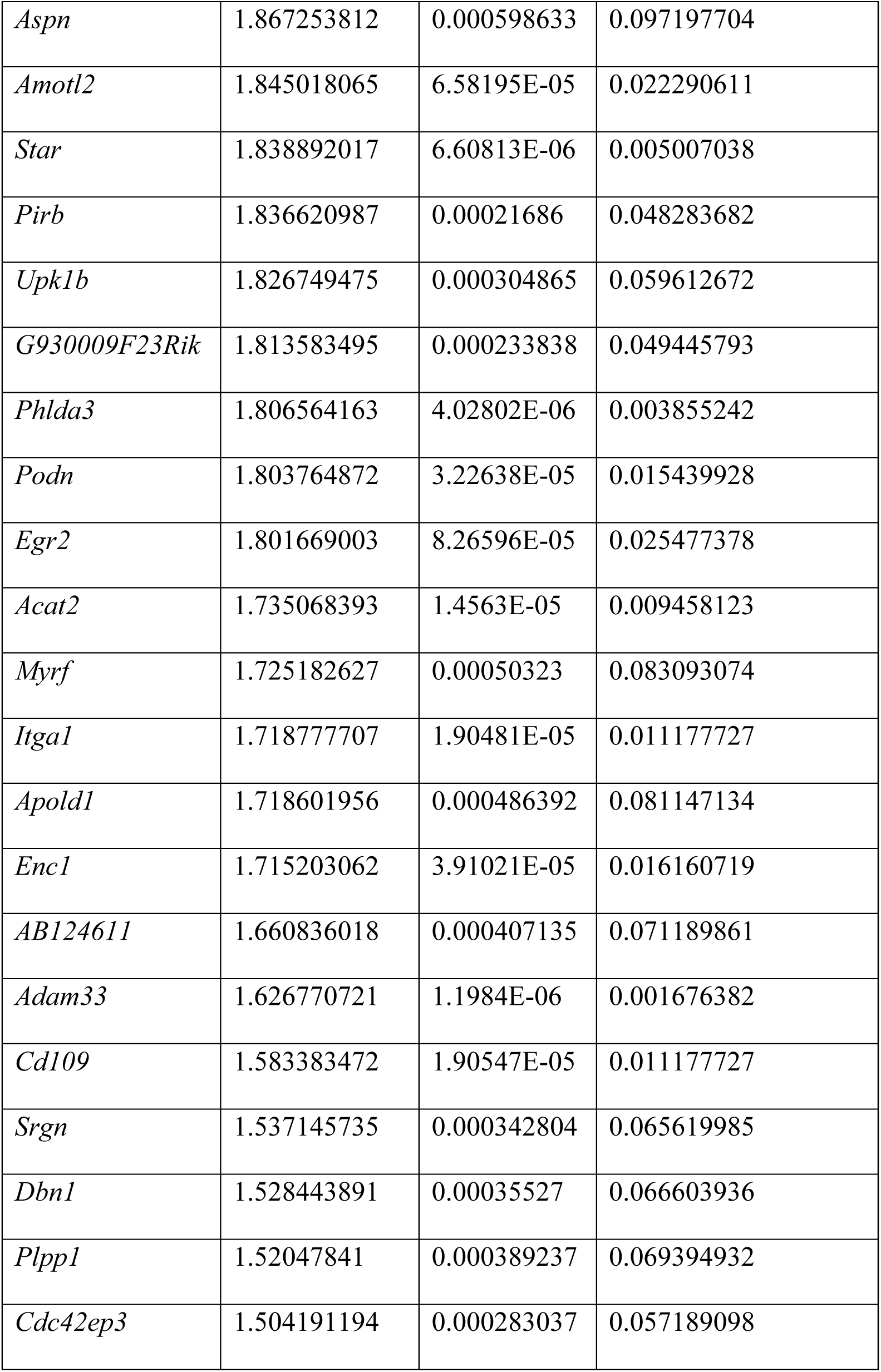
Top up-regulated genes in *Ccna2^d/d^* dpc 0.5 uteri.

**Supplemental Table 2.** Top down-regulated genes in *Ccna2^d/d^* dpc 0.5 uteri.

| **<u>Gene symbol</u>** | **Fold change** | **P-value** | **FDR** |
| --- | --- | --- | --- |
| *Dcc* | -6.92122 | 2.09E-05 | 0.011542 |
| *Ovgp1* | -4.55667 | 0.000209 | 0.048117 |
| *C4bp* | -3.95598 | 0.000265 | 0.055363 |
| *Gm19181* | -3.58923 | 2.53E-06 | 0.00288 |
| *Klf15* | -2.89858 | 2.08E-05 | 0.011542 |
| *Nlrp6* | -2.84518 | 6.92E-05 | 0.02248 |
| *Arl4d* | -2.81501 | 0.000115 | 0.031115 |
| *Lrrc75b* | -2.64622 | 3.68E-05 | 0.016027 |
| *Kcnc1* | -2.47186 | 4.91E-05 | 0.018597 |
| *Cacna1e* | -2.34272 | 0.000117 | 0.031305 |
| *Col6a6* | -2.22427 | 1.51E-07 | 0.000344 |
| *Zfp599* | -2.09059 | 1.33E-05 | 0.008953 |
| *Agmo* | -2.05983 | 0.000141 | 0.035739 |
| *Sned1* | -2.05738 | 0.000618 | 0.09864 |
| *Pthlh* | -2.01312 | 0.00022 | 0.048284 |
| *Gria4* | -1.97096 | 0.000299 | 0.059467 |
| *Gm48898* | -1.96663 | 6.36E-06 | 0.005007 |
| *Homer2* | -1.84314 | 7.54E-06 | 0.005484 |
| *Trpv4* | -1.7217 | 4.59E-05 | 0.017769 |

**Supplemental Table 3.** Antibodies used for immunohistochemical localization.

| **Antibody** | **Catalog number** | **Dilution** | **Vendor** |
| --- | --- | --- | --- |
| Cyclin A2 (CCNA2) | Ab181591 | 1:500 | Abcam<br>(Cambridge, MA) |
| Ki67 | Ab16667 | 1:100 | Abcam |
| Estrogen receptor-alpha<br>(ERα) | Ab32063 | 1:2500 | Abcam |
| Progesterone receptor A/B<br>(PGRA/B) | 8757 | 1:1000 | Cell Signaling Technology<br>(Danvers, MA) |

**Supplemental Table 4.**
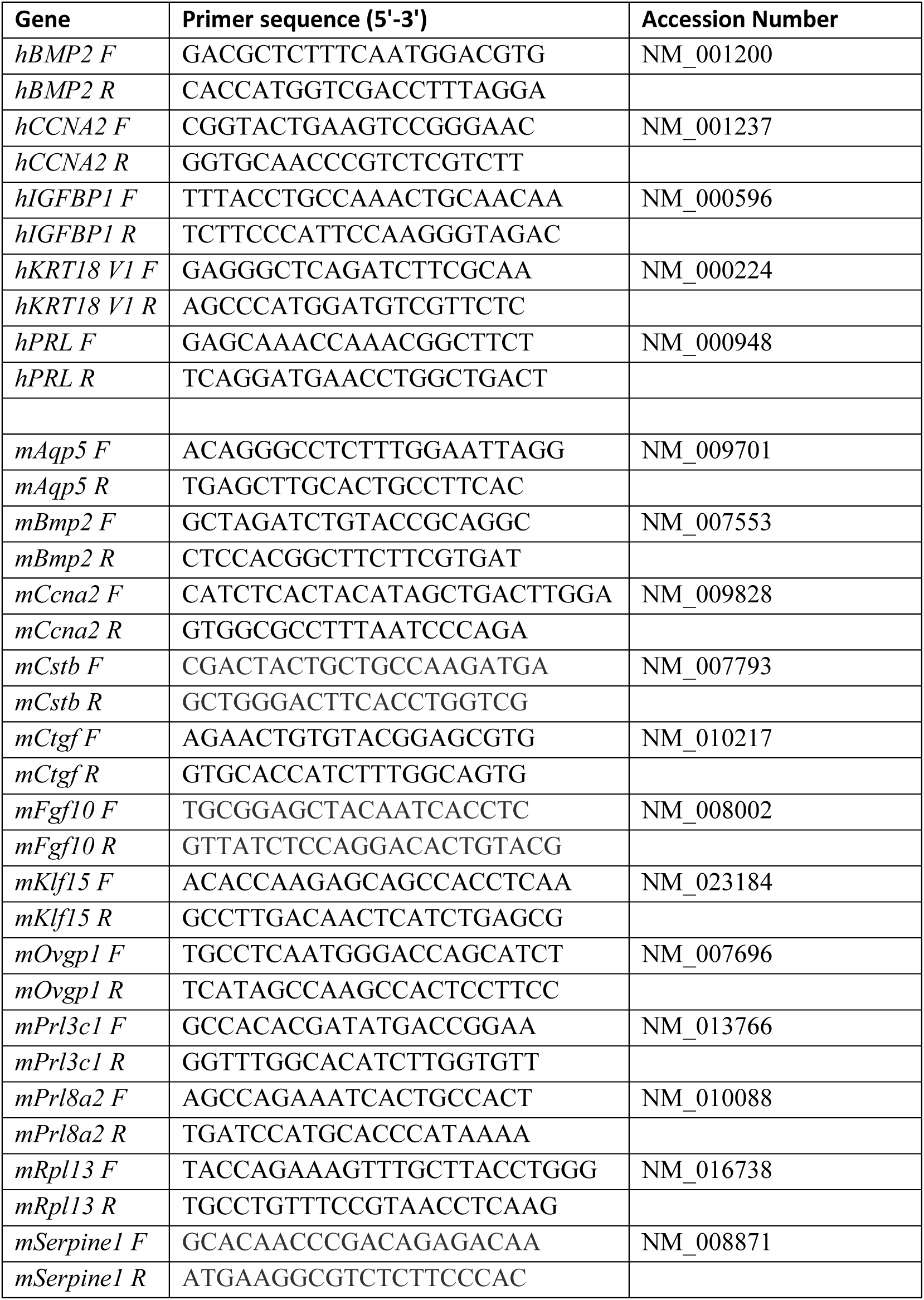
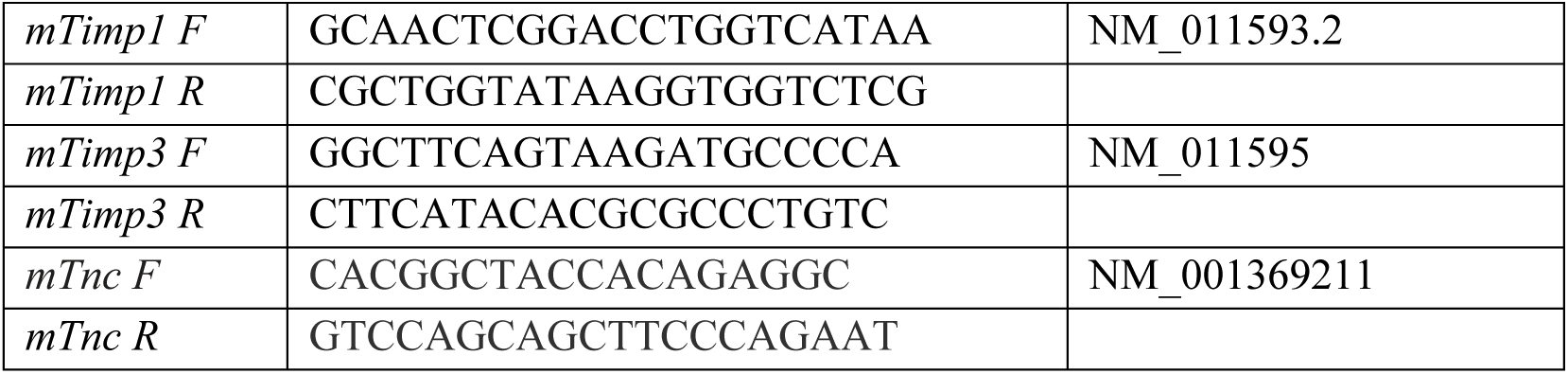
Human (h) and mouse (m) primers used for qRT-PCR.

